# Compensatory Creatine Secretion Protects Against Drug-Induced Liver Injury via Macrophage PHB1-K128 Desuccinylation: An NDRG1-Driven Metabolic – Immune Axis in APAP Hepatotoxicity

**DOI:** 10.64898/2026.11.15.766767

**Authors:** Yujun Luo, Mingkai Li, Yiming Lei, Jiayu Huang, Yan Chen, Hui Yuan, Yongcheng Xu, Weichun Lin, Hongsheng Yu, Huixin Chen, Felix W Leung, Hao Chen

**Author notes:** Contributing equally (Yujun Luo, Mingkai Li, Yiming Lei, Jiayu Huang).

## Abstract

**Background and aims:** Effective therapeutic interventions for Drug induced liver injury (DILI) remain limited. N-myc downstream regulated gene-1 (*NDRG1*), a stress-responsive proteins, has an incompletely defined role in acetaminophen-induced liver injury.

**Method:** Wild-type (WT), *NDRG1*^Hep-flox^, Hepatocyte-specific *NDRG1* conditional knockout mice (*NDRG1*^Hep-KO^) were subjected to APAP-induced injury modeling. Liver and blood sample from patient were analyzed. Liver sample of mice subsequently underwent RNA sequencing (RNA-Seq), untargeted metabolomic profiling, single-cell sequencing, succinylation sequencing. For *in vitro* studies, AML-12 hepatocytes, RAW264.7 macrophages, and bone marrow-derived macrophages (BMDMs), human Monocyte-Derived Macrophages, human hepatocyte organoids were utilized in experiments.

**Result:** *NDRG1* was upregulated in mouse and human DILI livers. *NDRG1* conditional knockout significantly exacerbates DILI. Mechanistically, *NDRG1* inhibited GAMT ubiquitination to enhance creatine production, and hepatocyte-derived creatine attenuated M1 polarization of liver macrophages by suppressing PHB1 succinylation at lysine 128. CXCL2⁺ macrophages were identified as a major subpopulation regulated by NDRG1.

**Conclusion:** Hepatocyte *NDRG1* protects against APAP-DILI by driving GAMT-dependent creatine production and PHB1 K128 succinylation-dependent macrophage reprogramming. Serum creatine may serve as a prognostic biomarker, and the *NDRG1*– creatine axis is a candidate therapeutic target.

**Bullet Points:**

1. Hepatocyte-specific knockout of *NDRG1* significantly aggravates DILI.
2. *NDRG1* in hepatocytes inhibits DILI M1 polarization of macrophages by regulating creatine synthase and thus creatine production.
3. *NDRG1*-mediated creatine secreted by hepatocytes by promoting PHB1 desuccinylation in macrophage.

**Graphic abstract:** NDRG1-regulated hepatocyte creatine inhibits APAP-induced M1 macrophage polarization via desuccinylation of PHB1 at K128

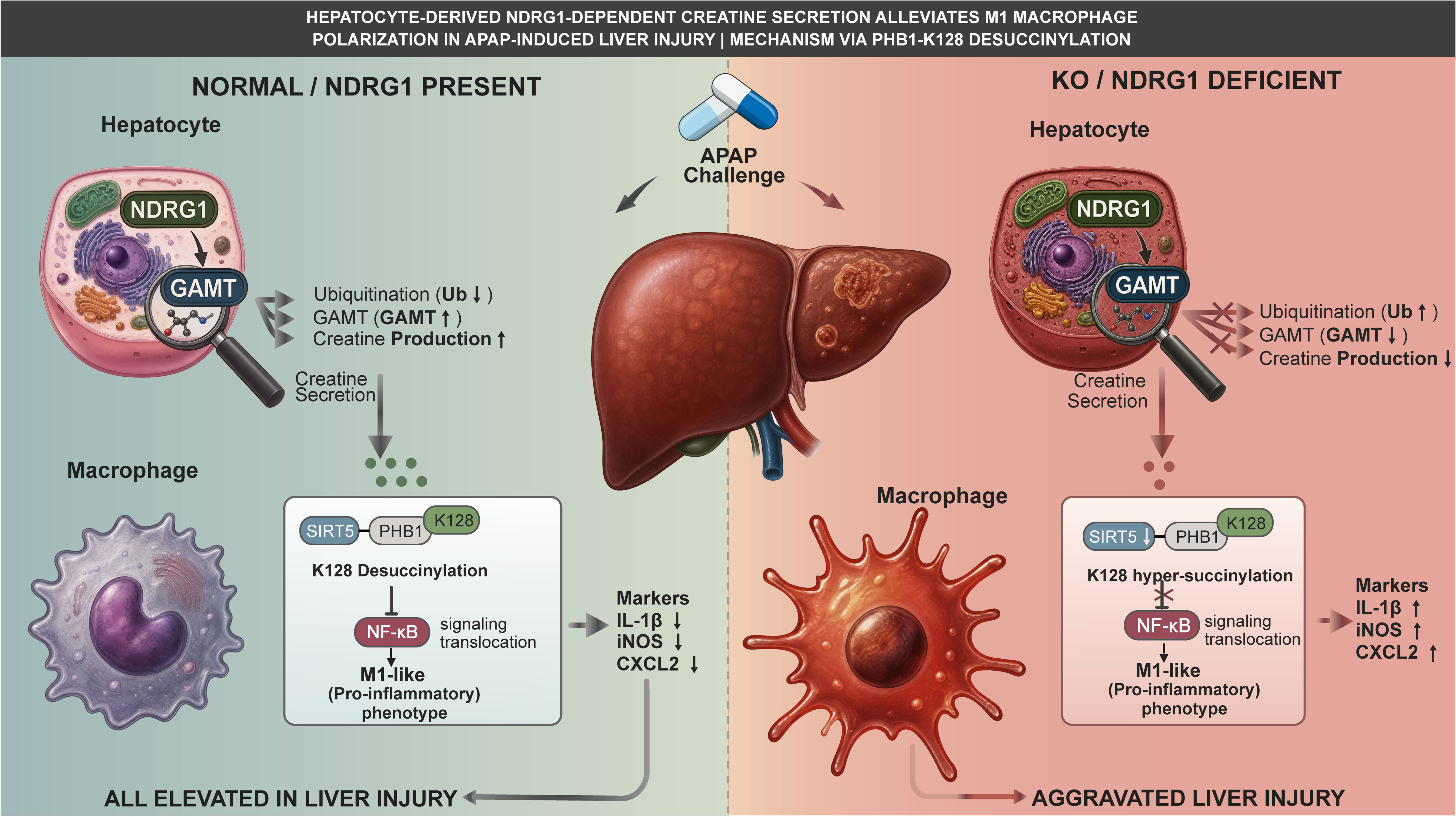

## 1. Introduction

Drug-Induced Liver Injury (DILI) refers to adverse effects on liver structure and function caused by medications, with clinical manifestations ranging from mild elevation of liver enzymes to severe hepatic dysfunction, including acute liver failure. A 2019 retrospective study estimated the incidence of DILI in China to be approximately 23.8/100, 000, with an increasing annual trend(Shen, et al., 2019). Acetaminophen (APAP), an FDA-approved over-the-counter medication for pain relief and fever reduction, has seen significantly increased usage during the COVID-19 pandemic due to the surge in febrile patients. However, prolonged or excessive APAP intake is a well-documented cause of DILI. Despite extensive research on APAP-induced hepatotoxicity, no more effective therapies for APAP-related DILI have been established. Thus, the development of novel treatments for APAP-induced DILI is imperative.

Recent studies have highlighted the pivotal role of metabolic reprogramming in curing diseases. Creatine is an amino acid containing high-energy phosphate bonds in humans. Its biosynthesis critically depends on two key enzymes: L-arginine:glycine amidinotransferase (AGAT) and guanidinoacetate N-methyltransferase (GAMT)(Barcelos, et al., 2016). Initially, creatine was primarily used to enhance athletic performance and increase muscle mass(Wang, et al., 2018). Then, subsequent research has revealed its therapeutic potential in hepatic encephalopathy induced by liver failure, fatty liver disease, doxorubicin-induced liver injury(Deminice, et al., 2011; Aljobaily, et al., 2020). However, role of creatine in APAP-induced liver injury remains unclear.

In recent years, accumulating evidence has demonstrated that macrophage polarization is closely associated with the initiation and progression of various diseases, such as acute liver injury(Peng, et al., 2019; Liu, et al., 2021). M1 macrophages produce large amounts of pro-inflammatory cytokines, primarily participating in acute immune responses and antimicrobial defense while M2 macrophages are involved in parasite clearance and tissue repair(Wang, et al., 2021). In APAP-induced liver Injury, hyperlipidemia aggravates liver damage by enhancing M1 polarization(Wang, et al., 2019). Conversely, human stem cells mitigate APAP-induced injury by promoting M2 polarization via the cGAS-STING pathway(Hua, et al., 2019). Type I interferon signaling modulates disease progression of DILI by influencing macrophage polarization(Song, et al., 2023). Notably, The N-myc downstream regulated gene 1 (*NDRG1*) plays a multifaceted role in cellular stress responses, differentiation, proliferation, and inflammation(Larkin, et al., 2014; Yuan, et al., 2025). While its involvement in vascular inflammation and respiratory diseases has been documented, its function in acetaminophen (APAP)-induced hepatocyte injury remains less understood(Zhang, et al., 2025).

Protein succinylation is a post-translational modification that transfers a succinyl group to lysine residues, regulating protein function and gene transcription. In the liver, succinylation is dynamically controlled by succinyltransferases (e.g., KAT2A, HAT1) and desuccinylases (primarily SIRT5 and SIRT7)(Liu, et al., 2024). SIRT5 is the key desuccinylase that removes succinyl groups to maintain hepatic metabolic homeostasis. Under various liver injury conditions, SIRT5 is often downregulated, leading to hyper-succinylation of its substrates and promoting disease progression. In APAP-induced acute liver injury, reduced SIRT5 impairs desuccinylation of ALDH2 at K385, exacerbating oxidative stress and inflammation(Yu, et al., 2024). In metabolic dysfunction-associated steatotic liver disease (MASLD), free fatty acids trigger SIRT5 degradation via TRIM21 upregulation, causing hyper-succinylation of metabolic and inflammatory proteins that aggravate glucolipid disorders and chronic inflammation(Xiao, et al., 2026). Collectively, targeting desuccinylases such as SIRT5 holds therapeutic potential.

This study focuses on the effect of NDRG1 on APAP-induced liver injury and the possible mechanism.

## 2. Materials and methods

### 2.1 *In vivo* study

Hepatocyte-specific *NDRG1* conditional knockout (*NDRG1*-cKO) or (*NDRG1*^Hep-KO^) mice were constructed by GemPharmatech Co., Ltd.

Animal model was performed according to previous study(Luo, et al., 2024). Briefly, mice were maintained under sterile conditions with ad libitum access to water and standard chow, under a controlled light-dark cycle. Fasting was initiated for ≥8 hours prior to experiments. On the day of the experiment, mice (8-10 weeks male 20g) received an intraperitoneal (i.p.) injection of 400 mg/kg acetaminophen (APAP) or PBS (every group has at least 3 mice). After 6 hours, mice were euthanized for tissue collection. For the survival analysis, mice were administered a higher dose of 500 mg/kg APAP (i.p.) and monitored for mortality over 72 hours. For survival analysis, 9 mice were in *NDRG1*^Hep-flox^ group while 10 mice were in *NDRG1*^Hep-KO^ group.

Mice with disabilities or malformations, females, anEMBOd those with other evident diseases were excluded from the study. Eight-week-old male mice weighing 20 g were identified as the optimal subjects for the experiment.

Randomisation was used to allocate experimental units to control and treatment groups.

Approximately 300 mice were used in the total experiment.

### 2.2 *In vitro* study

AML-12 and RAW 264.7 cell lines were obtained from ATCC. DMEM with 10%FBS (Gibco) were used to culture both cells.

BMDM were isolated from C57BL/6 mice. The isolation method was based on previous study. For conditioned medium study, AML-12 were treated with APAP or not for 24h. Then the supernatant was collected as CM. The BMDM were then cultured with CM for at less 24h.

Details are shown in the supplementary materials.

## 3. Result

### 3.1 Hepatocytic *NDRG1* is involved in acetaminophen (APAP)-induced liver injury

We first performed single-cell sequencing analysis on mouse livers. UMAP and dot plot analyses revealed nine distinct cell clusters in the mouse liver. Among these, the proportion of hepatocytes was significantly reduced in the APAP group compared with the control group (Figure 1 A, B, C). Further analysis indicated that NDRG1 mRNA expression in hepatocytes was significantly increased after APAP-induced injury, whereas its expression in hepatic macrophages showed no significant change (Figure 1 D, E). The mRNA and protein change of NDRG1 in hepatocyte were then confirmed by qPCR and western blot (Figure 1F, G, *P* < 0.05). Consistent with the single-cell sequencing results, fluorescence co-localization experiments revealed that *NDRG1* is predominantly localized in hepatocytes (Figure 1H). To further validate the single-cell sequencing results, hepatocytes were isolated and subjected to RNA transcriptome sequencing. The mRNA levels of *NDRG1* in hepatocytes were found to be significantly elevated following APAP-induced injury (Figure 1I, J). Moreover, the single-cell atlas of APAP-associated liver failure revealed that *NDRG1* is predominantly expressed in hepatocytes and is markedly elevated in hepatocytes from patients compared with those from healthy controls, consistent with previous findings in animal models. (Figure 1 K, L, M). Consistent with the above findings, NDRG1 expression was significantly increased in patients with drug-induced liver failure compared with healthy controls.(Figure 1N). Taken together, these findings indicate that *NDRG1* may participate in the pathogenesis of APAP-induced liver injury.

**Figure 1.**
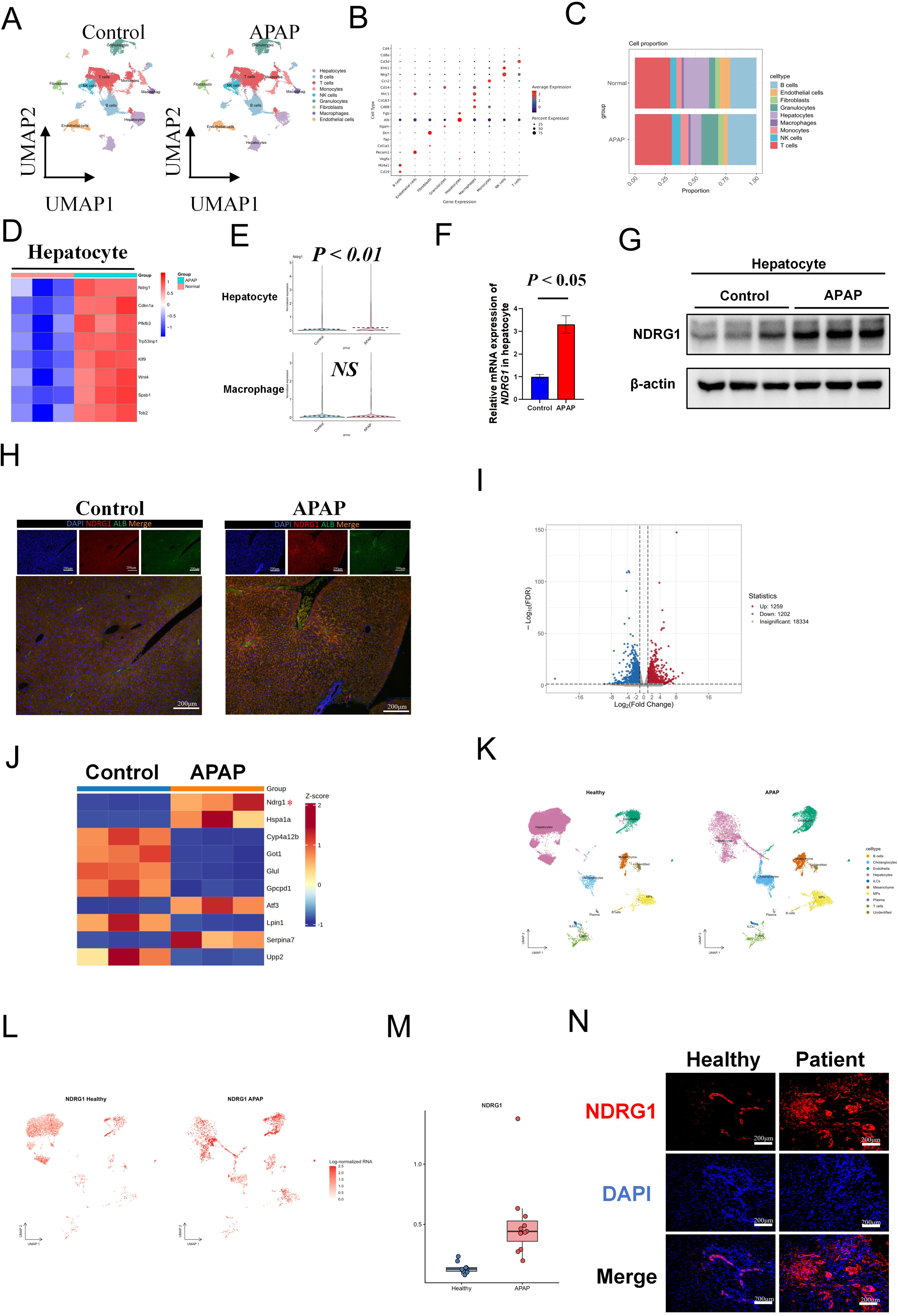
Hepatocytic NDRG1 is involved in acetaminophen (APAP)-induced liver injury (A) UMAP plot of single-cell transcriptomes of GEO228305. Control vs apap. (B) Cell-type annotation validation plot. (C) Cell proportion between normal and apap. (D) Heatmap of hepatocyte between normal and apap group. (E)Violin plots of gene expression of *NDRG1* in hepatocyte and macrophage. (F) mRNA expression of *NDRG1* in hepatocyte of mice. *P* < 0.05. (G)Protein expression of *NDRG1* in hepatocyte of mice. (H) Immunofluorescence colocalization of NDRG1 (red) and ALB (green). (I)(J) Volcano plot and heatmap of RNA-seq data from mouse liver hepatocytes. (K) Single-cell RNA sequencing data from GSE223581 were used to generate UMAP plots between healthy human liver and APAP-induced acute liver failure. (L) Feature plots illustrate the expression distribution of *NDRG1* in hepatocyte. (M) Box plot of NDRG1 expression. *P* < 0.05 (Wilcoxon). (N) Immunofluorescence staining showing hepatic NDRG1 expression in healthy controls and patients with drug-induced liver failure.

### 3.2 Deficiency of *NDRG1* in hepatocyte aggravates APAP-induced liver injury

To further investigate the role of NDRG1 in APAP-induced liver injury, we generated NDRG1^Hep-KO^ mice (Figure 2A). Conditional deletion of *NDRG1* in hepatocytes significantly exacerbated APAP-induced liver necrosis, transaminase elevation, inflammation, and oxidative stress (Figure 2B, C, D, E). In addition, TUNEL and MPO staining demonstrated that hepatocyte-specific *NDRG1* knockout markedly promoted hepatocyte apoptosis, neutrophil infiltration and increased the proportion of macrophages (Figure 2F, G, H *, vs *NDRG1*^Hep-flox^ + APAP. #, vs *NDRG1*^Hep-flox^. *P* < 0.05). Furthermore, survival analysis revealed a markedly reduced survival rate in *NDRG1*^Hep-KO^ mice compared to NDRG1^Hep-flox^ controls following high-dose APAP exposure (Figure 2I, *P* < 0.05). Collectively, these findings indicate that *NDRG1* in hepatocyte may serve as a protective factor against APAP-induced hepatotoxicity.

**Figure 2.**
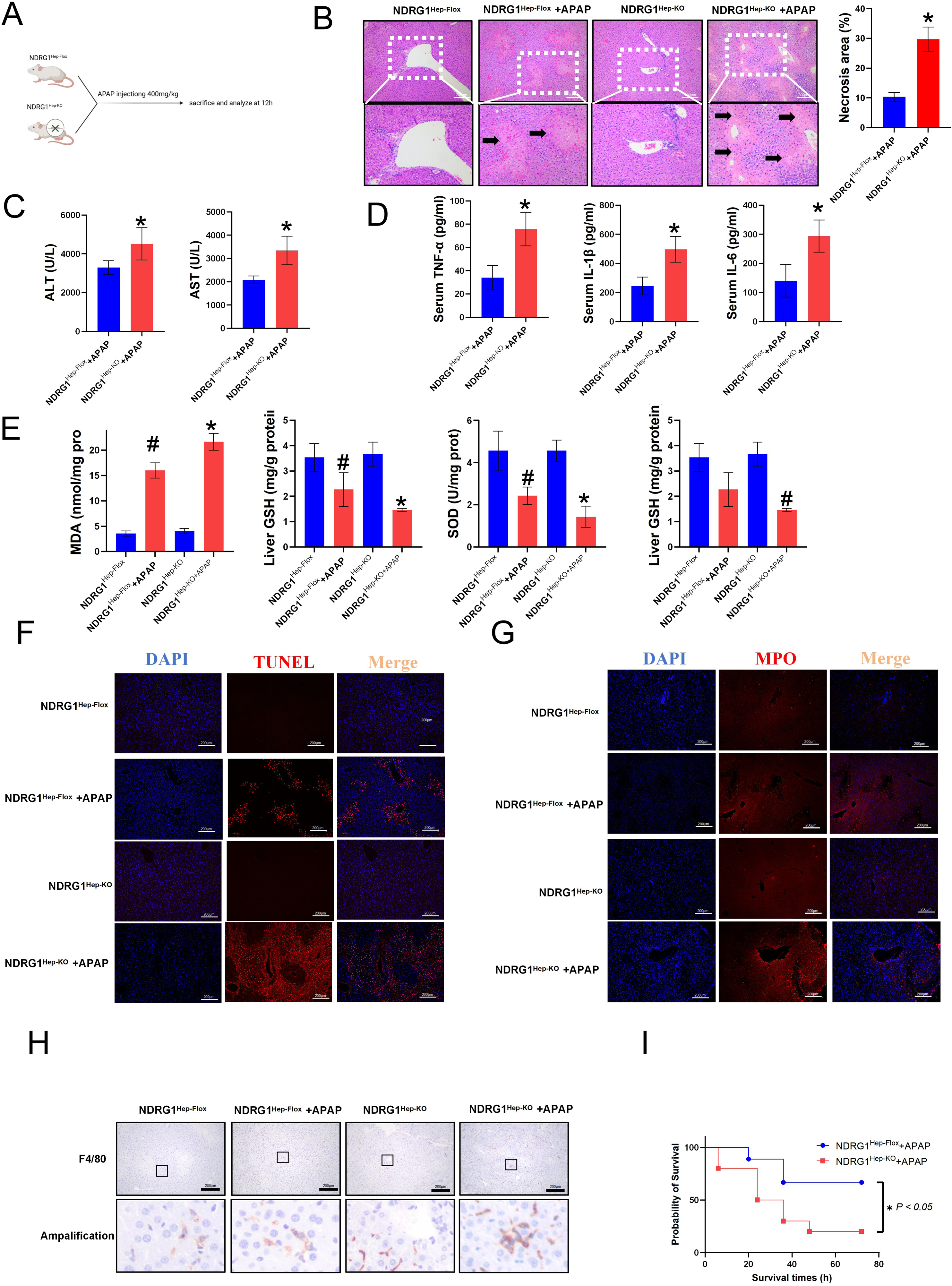
Deficiency of NDRG1 in hepatocyte aggravates APAP-induced liver injury. (A) Experimental design.(B) Representative H&E staining of mouse liver sections and quantification of necrotic area. *P* < 0.05 versus *NDRG1*^Hep-flox^+APAP; n = 3 per group.(C, D) Serum levels of AST, ALT, TNF-α, IL-1β, and IL-6 in mice. *P* < 0.05 versus *NDRG1*^Hep-flox^+APAP; n = 3.(E) Hepatic contents of MDA, GSH, SOD, and GPX. #, versus *NDRG1*^Hep-flox^; * versus *NDRG1*^Hep-flox^+APAP, *P* < 0.05, n = 3. (F, G) Immunofluorescence staining for TUNEL and MPO. *, versus *NDRG1*^Hep-flox^. #, versus *NDRG1*^Hep-flox^+APAP *P* < 0.05. n = 3. (H) Immunohistochemical staining for F4/80 in the liver. (I) Kaplan–Meier survival analysis. *P* < 0.05.

### 3.3 *NDRG1* deficiency exacerbates M1 polarization of hepatic macrophages in APAP-induced liver injury

To further explore the potential mechanism underlying the protective role of *NDRG1*, we performed single-cell sequencing on livers of *NDRG1* cKO mice and *NDRG1*^Hep-flox^ mice following APAP-induced injury. UMAP plots and cell type annotation based on defined marker genes revealed ten distinct cell populations in the liver (Figure 3A, B). The proportion of hepatocytes was markedly decreased in *NDRG1*^Hep-KO^ mice, whereas the proportion of macrophages was significantly increased, suggesting an important role for macrophages in this process (Figure 3C). Further analysis of M1 polarization markers in macrophages showed that the expression of NOS2, TNF-α, IL-1β and other molecules was significantly elevated in hepatocyte-specific *NDRG1*^Hep-KO^ mice, indicating that macrophage M1 polarization may play a key role (Figure 3D).

**Figure 3.**
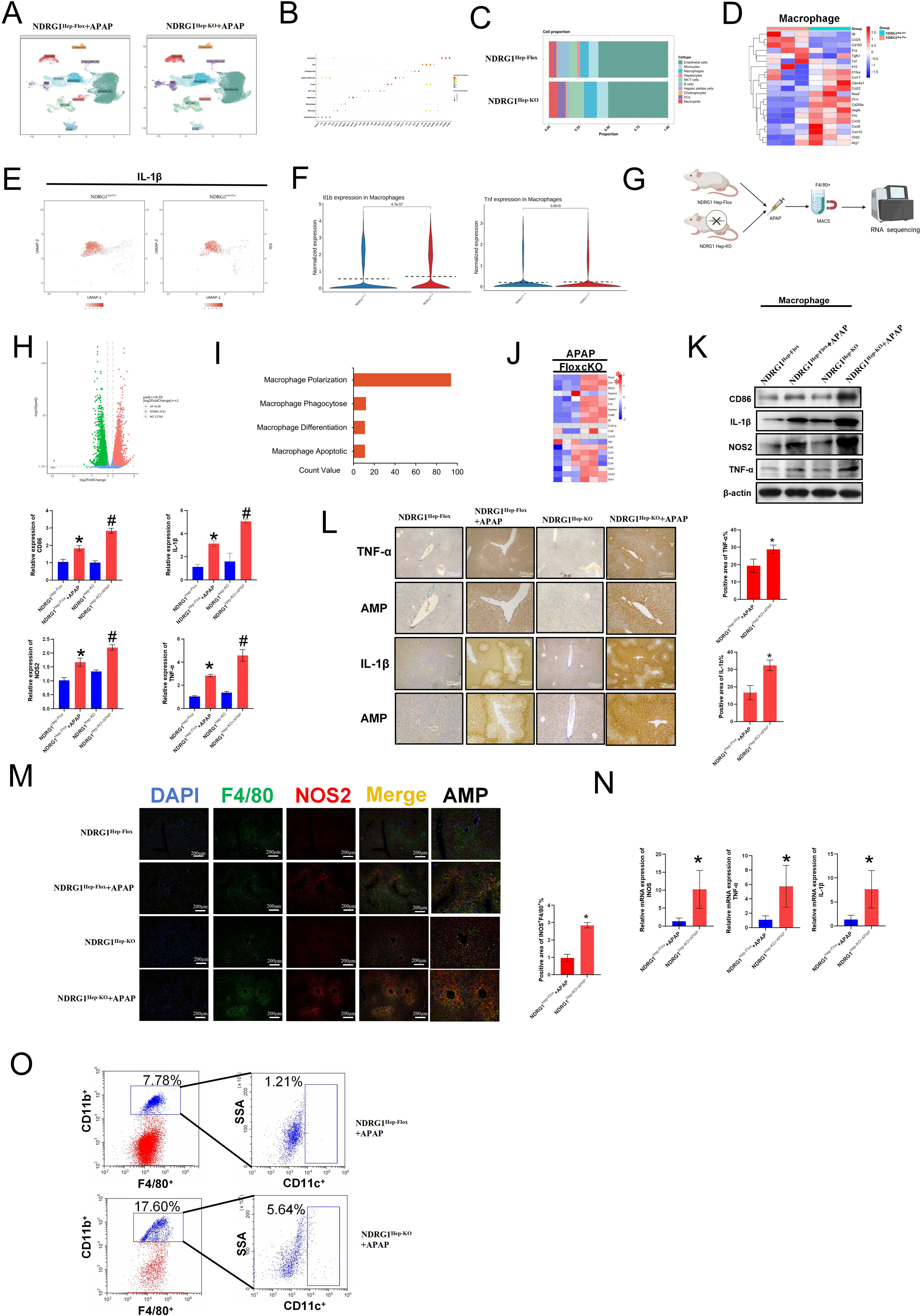
*NDRG1* deficiency exacerbates M1 polarization of hepatic macrophages in APAP-induced liver injury *in vivo*. (A) UMAP plot of single-cell transcriptomes. *NDRG1*^Hep-flox^+APAP Vs *NDRG1*^Hep-KO^+APAP. (B) Cell-type annotation validation plot. (C)) Cell proportion between *NDRG1*^Hep-flox^+APAP and *NDRG1*^Hep-KO^+APAP. (D) Single-cell RNA-seq heatmap showing differentially expressed genes in macrophages between *NDRG1*^Hep-flox^+APAP and *NDRG1*^Hep-KO^+APAP groups. (E) Feature plot of IL-1β expression.(F) Violin plots comparing IL-1β and TNF expression between *NDRG1*^Hep-flox^+APAP and *NDRG1*^Hep-KO^+APAP groups. *P* < 0.05. (G) Schematic diagram of the experimental design. (H) Volcano plot of bulk RNA-seq data (*NDRG1*^Hep-flox^+APAP vs. *NDRG1*^Hep-KO^+APAP). (I) Gene Set Enrichment Analysis (GSEA) plots. (J) Heatmap of bulk RNA-seq expression profiles. (K) Western blot analysis of proteins in mouse liver macrophages. * versus *NDRG1*^Hep-flox^, *P* < 0.05. # versus *NDRG1*^Hep-flox^+APAP, *P* < 0.05. (L, M) Immunohistochemistry and immunofluorescence staining of mouse liver sections.* versus *NDRG1*^Hep-flox^+APAP, *P* < 0.05. (N) mRNA expression levels in mouse liver macrophages. * vs *NDRG1*^Hep-flox^+APAP, *P* < 0.05. (O) Flow cytometric analysis of whole liver cells from mice.

To further investigate the role of macrophages, we isolated macrophages from mouse liver tissues using F4/80 magnetic beads and subjected them to transcriptome sequencing (Figure 3G, H). GSEA indicated that the increased macrophages in *NDRG1*^Hep-KO^ mice were primarily associated with polarization (Figure 3I). A heatmap demonstrated that M1 polarization markers, including NOS2 and IL-1β, were significantly upregulated in *NDRG1*^Hep-KO^ mice (Figure 3J). Furthermore, qPCR, western blotting, immunohistochemistry, and immunofluorescence double staining consistently confirmed that the degree of M1 polarization was significantly higher in *NDRG1*^Hep-KO^ mice than in *NDRG1*^Hep-flox^ mice (Figure 3J, K, L, M, N). In addition, flow cytometry analysis revealed a marked increase in F4/80⁺ CD11b⁺ CD11c⁺ triple-positive cells in the *NDRG1*^Hep-KO^ group (Figure 3O).

To further verify the role of M1 polarization in *NDRG1*-mediated protection, we collected supernatants from AML12 cells and co-cultured them with RAW264.7 macrophages (Supplementary Figure 1A). Conditioned medium from APAP-stimulated *NDRG1*-knockdown AML12 cells, compared with that from siNC-treated AML12 cells, markedly enhanced M1 polarization of RAW264.7 cells, as shown by increased iNOS, IL-1β, and TNF-α levels via western blotting (Supplementary Figure 1B. *P* < 0.05). Flow cytometry further revealed that this conditioned medium promoted the conversion of RAW264.7 cells into F4/80⁺ CD11b⁺ CD86⁺ triple-positive cells (Supplementary Figure 1C). Additionally, bone marrow-derived macrophages (BMDMs) were isolated and co-cultured with AML12-conditioned medium (Supplementary Figure1 D). In line with the above findings, conditioned medium from APAP-stimulated *NDRG1*-knockdown AML12 cells also facilitated the conversion of BMDMs into F4/80⁺ CD11b⁺ CD86⁺ triple-positive cells (Supplementary Figure 1E, F, G). Taken together, lack of *NDRG1* in hepatocytes promotes M1 polarization of macrophages *in vivo* and *in vitro*.

### 3.4. *NDRG1* deficiency reduces hepatic *GAMT* expression and creatine content *in vivo*

To elucidate the mechanisms by which *NDRG1* deficiency influences APAP-induced liver injury, we performed untargeted metabolomic sequencing on hepatocytes. The results revealed 255 significantly upregulated metabolites and 192 significantly downregulated metabolites (Figure 4A). KEGG enrichment analysis of these differentially abundant metabolites indicated that the changes were primarily enriched in the arginine and proline metabolism pathway (Figure 4B). Notably, creatine metabolism is closely associated with arginine metabolism (Figure 4C). Subsequent sequencing and validation experiments demonstrated a significant decrease in hepatic creatine content following *NDRG1* knockout (Figure 4D, E, *P* < 0.05). Therefore, we hypothesized that *NDRG1* may be critically involved in creatine metabolism.

**Figure 4.**
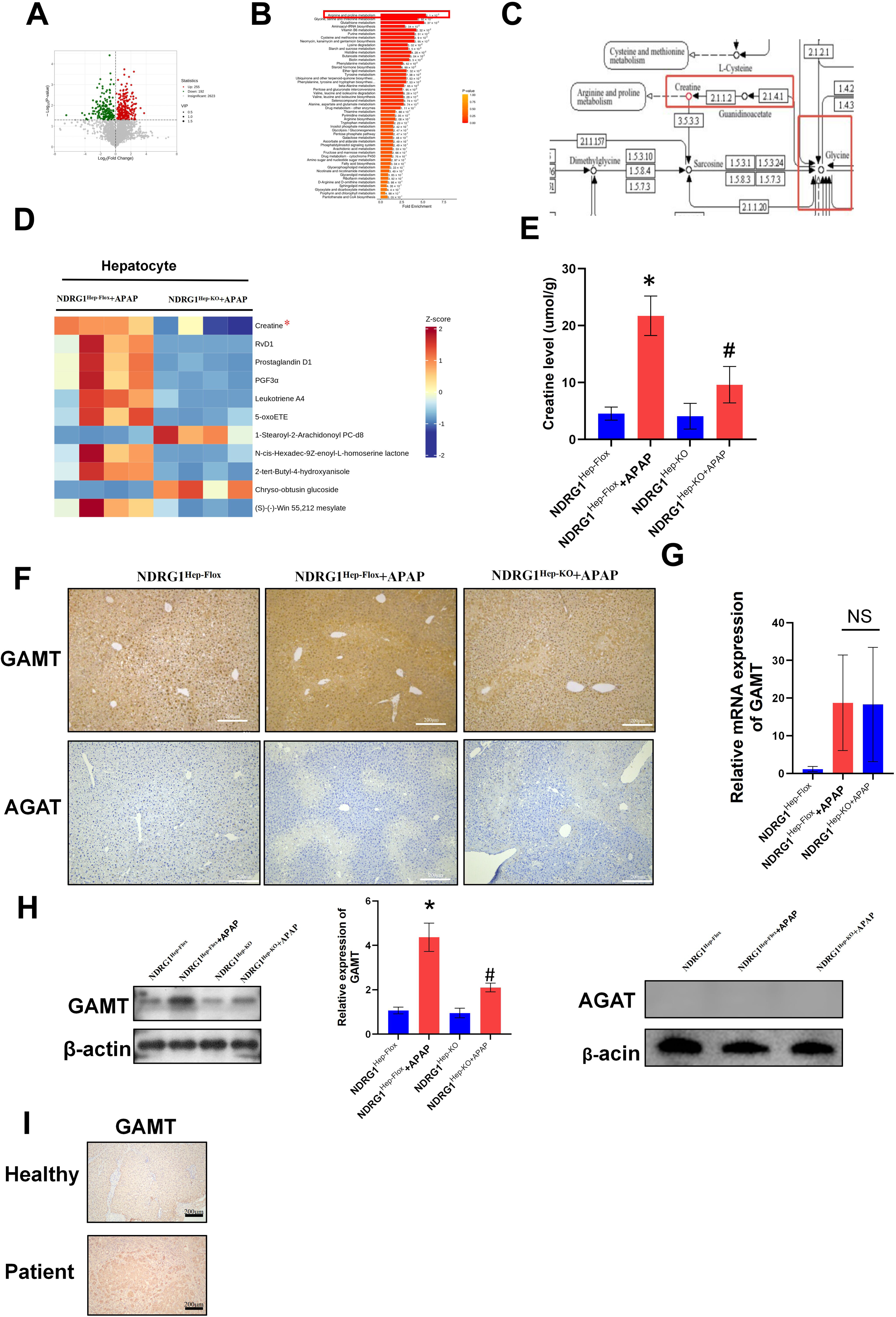
*NDRG1* deficiency reduces hepatic GAMT expression and creatine content *in vivo*. (A) Volcano plot shows the differential metabolites of hepatocyte between *NDRG1*^Hep-flox^ and *NDRG1*^Hep-KO^ mice challenged with APAP. (B) KEGG of the differential metabolites of hepatocyte between *NDRG1*^Hep-flox^ and *NDRG1*^Hep-KO^ mice challenged with APAP. (C) Schematic Diagram of the Metabolic Pathway. (D) Heat map of the differential metabolites of hepatocyte between *NDRG1*^Hep-flox^ and *NDRG1*^Hep-KO^ mice challenged with APAP. (E) Hepatic creatine level. * vs *NDRG1*^Hep-flox^ *P* < 0.05, # vs *NDRG1*^Hep-flox^ + APAP *P* < 0.05. n=3. (F) Immunohistochemical Staining of GAMT and AGAT in mouse liver. (G) RT-PCR of liver tissue. * vs *NDRG1*^Hep-flox^ + APAP *P* < 0.05. (H) Representative immunoblots of hepatocyte in liver tissue.* vs *NDRG1*^Hep-flox^ *P* < 0.05, # vs *NDRG1*^Hep-flox^ + APAP *P* < 0.05. n=3. (I) Representative immunohistochemistry images showing hepatic GAMT expression in healthy controls and patients with drug-induced liver failure.

Given that GAMT and AGAT are the rate-limiting enzymes for creatine synthesis, we conducted additional experiments. Western blot and immunohistochemical analyses revealed a significant decrease in GAMT protein levels in *NDRG1*^Hep-KO^ hepatocytes, while AGAT expression showed no significant difference (Figure 4F, H, *P* < 0.05). Following APAP-induced injury, GAMT mRNA levels were markedly elevated, whereas knockout of *NDRG1* in hepatocytes exerted no influence on GAMT mRNA levels (Figure 4G, *P* > 0.05). To further validate the clinical relevance of GAMT and creatine in patients with drug-induced liver injury, liver specimens were collected from patients with APAP-related drug-induced liver failure and compared with those from healthy controls. Consistent with the animal experiments, immunohistochemical analysis revealed that hepatic GAMT protein expression was markedly higher in patients with APAP-related liver failure than in healthy controls (Figure 4I).Previous study indicates that *GAMT* is primarily expressed in the liver, whereas AGAT is predominantly expressed in the kidneys.

Collectively, these experiments demonstrate that *NDRG1* knockout suppresses the expression of *GAMT* and consequently reduces the levels of its metabolic product, creatine.

### 3.5 *NDRG1* suppresses M1 polarization of hepatic macrophages by augmenting creatine synthesis in hepatocytes

Creatine is primarily synthesized in hepatocytes. To further investigate whether *NDRG1* influences macrophage M1 polarization by modulating hepatic creatine synthesis, we supplemented the CM with creatine (Figure 5A). Experimental results demonstrated that knockdown of *NDRG1* significantly reduced creatine levels in the supernatant of APAP-injured AML-12 (Figure 5B, *P* < 0.05). Exogenous creatine supplementation partially reversed the effect of *NDRG1* knockdown on M1 polarization in RAW 264.7 macrophages as evidenced by attenuation of the enhanced protein expression of TNF-α, Il-1β, and NOS2, reduced levels of the pro-inflammatory cytokines IL-1β, IL-6, and TNF-α in the supernatant (Figure 5C, D, *P* < 0.05). Furthermore, flow cytometry analysis revealed that exogenous creatine supplementation significantly reversed the *NDRG1* knockdown-induced increase in the proportion of F4/80⁺ CD11b⁺ CD86⁺ macrophages (Figure 5E). Subsequently, we found that overexpression of *NDRG1* in AML-12 hepatocytes subjected to APAP-induced injury significantly increased GAMT expression and elevated creatine levels in the supernatant (Figure 5G, *P* < 0.05). Next, the supernatant derived from *NDRG1*-overexpressing cells was applied to RAW 264.7 macrophages and compared to that from the vector control group. The results demonstrated that *NDRG1* overexpression significantly suppressed M1 polarization in RAW 264.7 cells, as evidenced by western blot (Figure 5F, *P* < 0.05). Moreover, we observed that overexpression of GAMT itself also elevated creatine levels and inhibited M1 macrophage polarization (Figure 5I, *P* < 0.05). Collectively, these findings indicate that *NDRG1* suppresses M1 polarization of hepatic macrophages by activating GAMT in hepatocytes, thereby augmenting creatine secretion.

**Figure 5.**
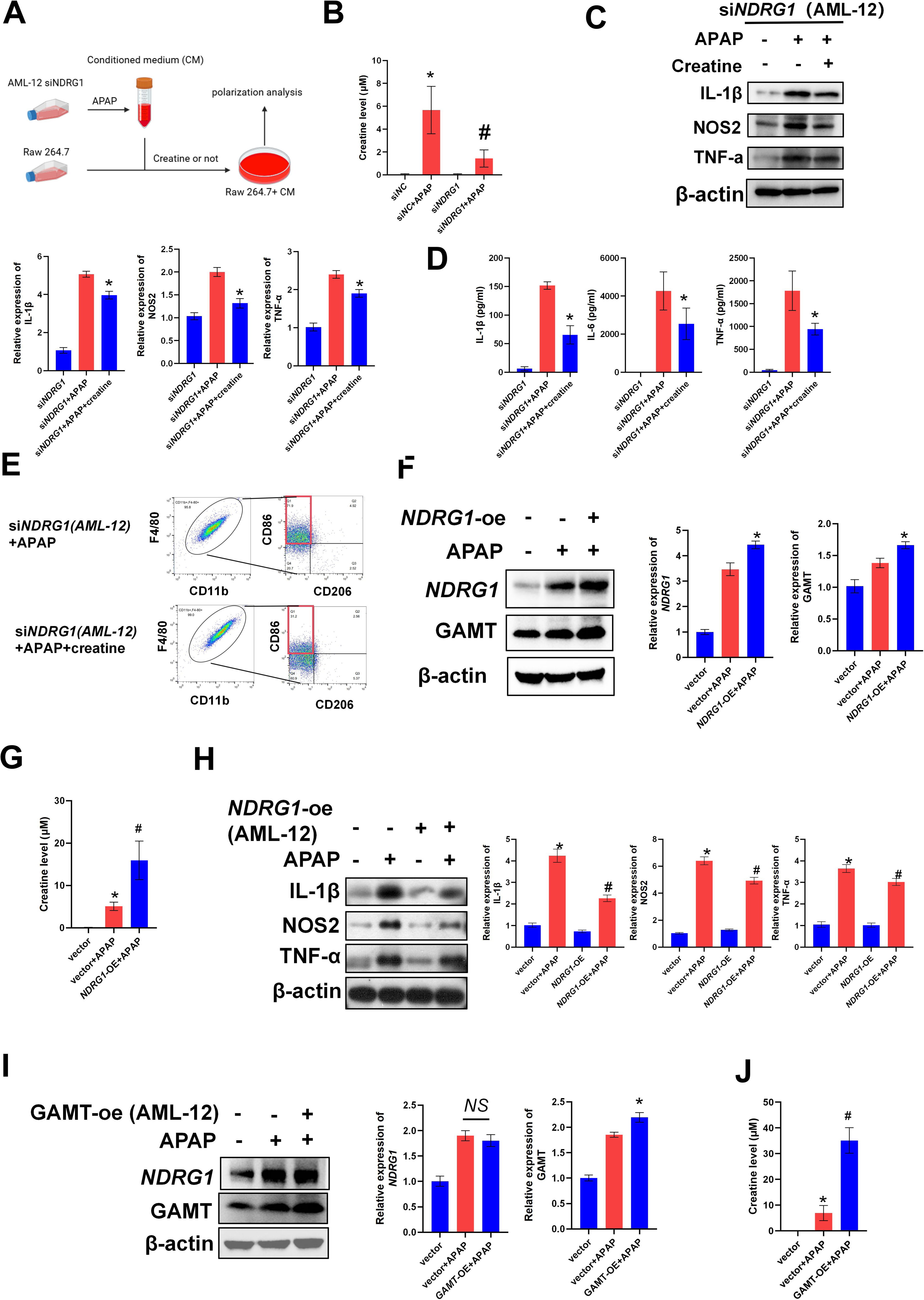
*NDRG1* suppresses M1 polarization of hepatic macrophages by augmenting creatine synthesis in hepatocytes. (A) Schematic diagram of *in vitro* study. RAW 264.7 cultured with conditioned medium were treated with or without creatine. (B) Creatine level in the supernatant of AML-12. * vs si*NC* + APAP *P* < 0.05, # vs si*NDRG1* + APAP *P* < 0.05. (C) Western blot analysis of M1 macrophage polarization marker in RAW 264.7.* vs si*NDRG1* + APAP *P* < 0.05. (D) Concentrations of inflammatory cytokines in supernatant of RAW 264.7.* vs si*NDRG1* + APAP *P* < 0.05. (E) Flow cytometric analysis of RAW 264.7 (F4/80^+^, CD86^+^, CD11b^+^). (F) Western blot of *GAMT* and *NDRG1* in AML-12. * vs Vector + APAP *P* < 0.05. (G) Creatine level in the supernatant of AML-12. * vs Vector *P* < 0.05, # vs Vector + APAP *P* < 0.05. (H) Western blot analysis of M1 macrophage polarization marker in RAW 264.7. * vs Vector + APAP *P* < 0.05, # vs *NDRG1*-OE + APAP *P* < 0.05. (I) Western blot of *GAMT* and *NDRG1* in AML-12. * vs Vector + APAP *P* < 0.05. (J) Creatine level in the supernatant of AML-12. * vs Vector *P* < 0.05, # vs Vector + APAP. Oe:overexpression.

### 3.6 Creatine supplementation significantly reversed the enhancement of macrophage M1 polarization induced by *NDRG1* conditional knockout

To verify that creatine deficiency is one of the key factors mediating the enhanced M1 polarization of macrophages induced by hepatocyte-specific *NDRG1* conditional knockout, mice were pretreated with creatine (Figure 6A). The results showed that exogenous creatine supplementation significantly reduced the area of hepatic necrosis and serum transaminase levels, and improved survival in APAP-treated mice (Figure 6B, C, D, *P* < 0.05, *, vs cKO + APAP). Transcriptome sequencing of macrophages isolated from mouse livers further showed that exogenous creatine supplementation suppressed M1 polarization, which was confirmed by immunohistochemistry, immunofluorescence, and western blotting, flowcytometry (Figure 6E, F, G, H, I, J *P* < 0.05). These findings suggest that creatine may be one of the key metabolites involved in the NDRG1-mediated suppression of M1 polarization.

**Figure 6.**
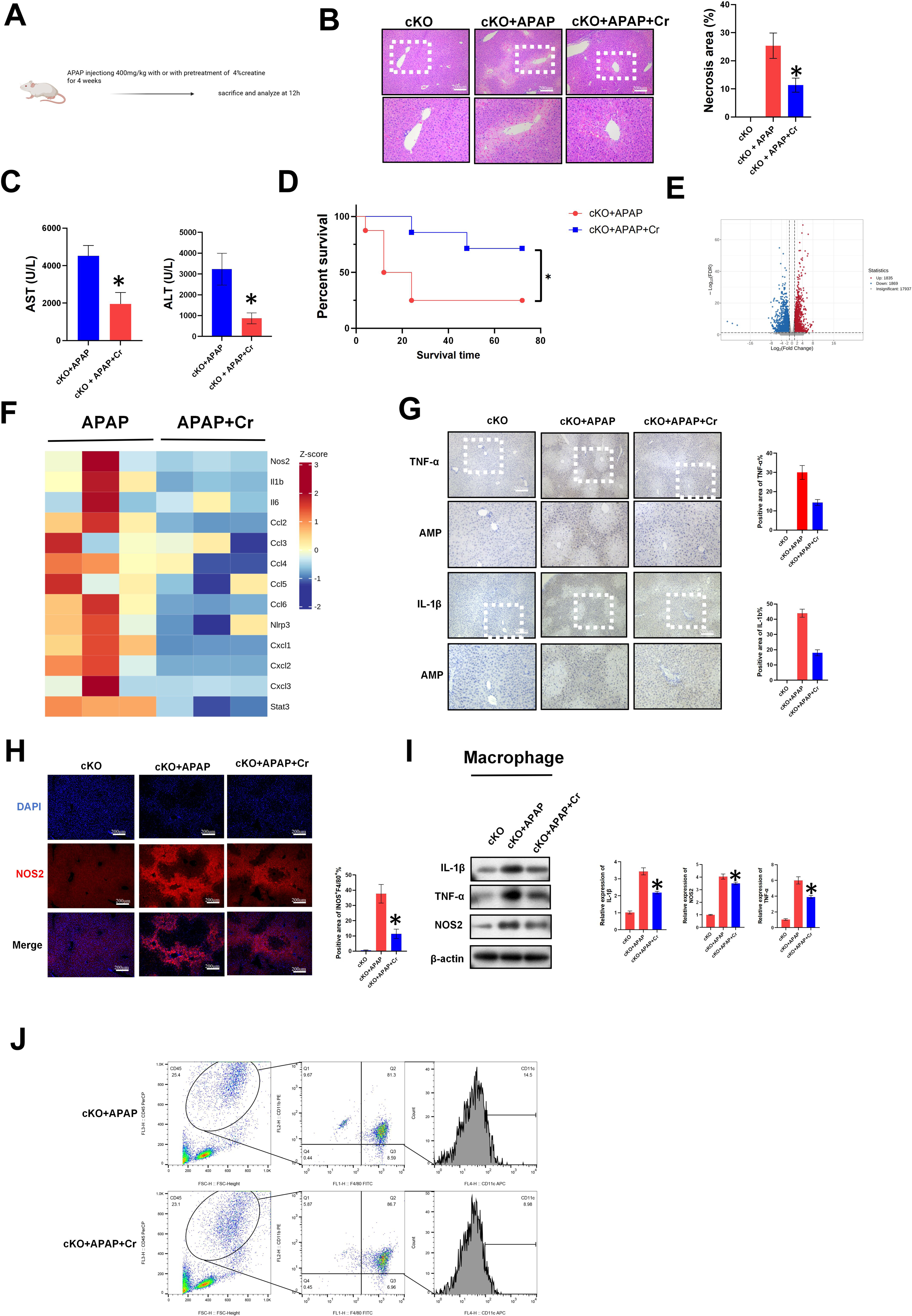
Creatine supplementation significantly reversed the enhancement of macrophage M1 polarization induced by *NDRG1* conditional knockout. (A) Experimental design (B) Mouse liver HE staining, necrotic area, vs *NDRG1*^Hep-KO^+APAP, *P* < 0.05, n=3, Cr=creatine. (C) Mouse serum AST and ALT levels。,vs *NDRG1*^Hep-KO^+APAP, P < 0.05, n=3(D) Mouse K-M survival analysis., vs *NDRG1*^Hep-KO^+APAP, *P* < 0.05. (E) Volcano plot in transcriptome sequencing of mouse primary macrophages。(F) Differential expression plot of mouse macrophage transcriptome. *NDRG1*^Hep-KO^+APAP vs *NDRG1*^Hep-KO^+APAP+creatine. (G) Immunohistochemical staining of TNF-α and IL-1β in mouse liver.,vs *NDRG1*^Hep-KO^+APAP, P < 0.05, n=3. (H) Immunofluorescence staining of NOS2 in mouse liver. (I) Western blot of mouse primary hepatic macrophages. *, vs *NDRG1*^Hep-KO^+APAP, *P* < 0.05, n=3.(J) Flow cytometry of mouse liver non-parenchymal cells.

### 3.7 *NDRG1* modulates *GAMT* expression through its critical amino acid residues Tyr-112, Ala-230, Ser-233, and Glu-238 and ubiquitination

Based on the established association between *NDRG1* and *GAMT*, we hypothesized a direct interaction between *NDRG1* and *GAMT*. Immunofluorescence co-staining revealed co-localization of *NDRG1* and *GAMT*, suggesting potential direct binding (Figure 7A). Co-immunoprecipitation (Co-IP) assays confirmed a direct physical interaction between *NDRG1* and *GAMT* (Figure 7B, C).

**Figure 7.**
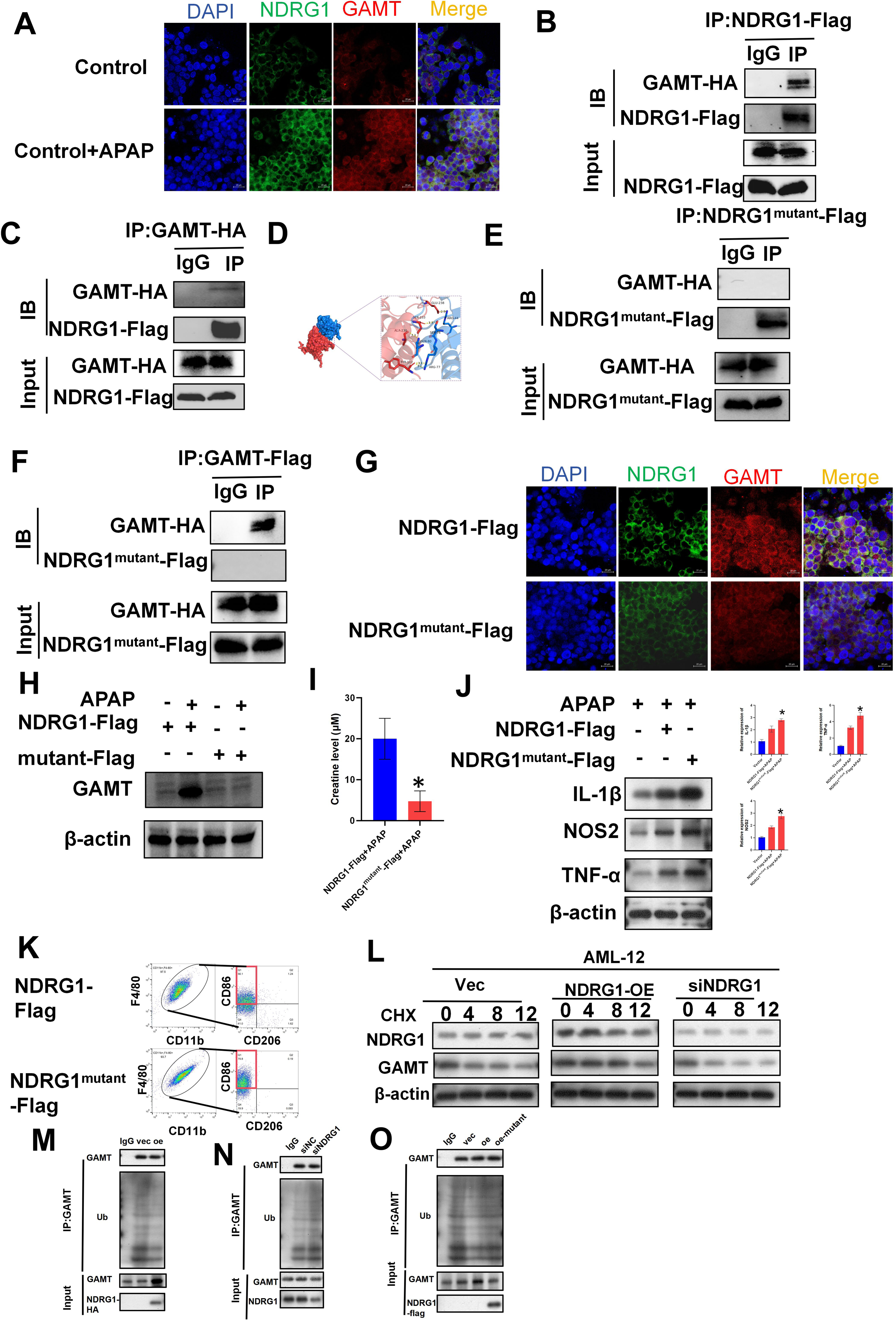
*NDRG1* modulates GAMT expression through its critical amino acid residues Tyr-112, Ala-230, Ser-233, and Glu-238 and ubiquitination. (A) Dual immunofluorescence staining of GAMT and NDRG1 in AML-12. (Red = GAMT, Green =NDRG1. (B) (C) Co-IP between exogenous NDRG1 and GAMT in AML-12 cells transfected with GAMT-HA and NDRG1-FLAG plasmids. (D) 3D structures of NDRG1/GAMT (E) (F) CoIP between exogenous NDRG1mutant and GAMT in AML-12 cells transfected with NDRG1mutant-FLAG and GAMT-HA plasmids. (G) Dual immunofluorescence staining of GAMT and NDRG1 in AML-12. (Red = GAMT, Green =NDRG1.(H) Western blot of GAMT in AML-12 transfected with the indicated plasmid. (I) Creatine level in the supernatant of AML-12 transfected with the indicated plasmid. * vs NDRG1-FLAG + APAP P < 0.05. (J) Western blot of M1 macrophage polarization markers in RAW 264.7 co-cultured with CM from AML-12 transfected with the indicated plasmid.* vs NDRG1-FLAG + APAP P < 0.05. (K) Flow cytometric analysis of RAW 264.7 (F4/80+, CD86+, CD11b+).(L) Cell synthesis inhibition assay, CHX inhibition time expressed in hours. (M)(N)(O) Mouse RAW264.7 cell co-IP assay, IP: *GAMT*.

Using UniProt-based binding site prediction, we identified putative interaction residues in *NDRG1* (Tyr-112, Ala-230, Ser-233, Glu-238) (Figure 7D). Consequently, we generated site-directed mutagenesis plasmids (Tyr112Ser, Ala230Asp, Ser233Arg, Glu238Asp) and transfected them into AML-12 hepatocytes, with wild-type (WT) *NDRG1* plasmid as control. Co-IP analysis demonstrated significantly weakened *NDRG1-GAMT* binding upon mutation of these residues, indicating that Tyr-112, Ala-230, Ser-233, and Glu-238 mediate the interaction (Figure 7E, F). This impaired binding was corroborated by diminished co-localization in immunofluorescence (Figure 7G). Concurrently, AML-12 cells expressing mutant *NDRG1* exhibited reduced *GAMT* expression and lower creatine content compared to WT-transfected controls (Figure 7H, I, *P* < 0.05). Functional assessment revealed that CM from mutant groups exacerbated M1 macrophage polarization in RAW 264.7 cells versus WT-CM, evidenced by western blot and flow cytometry (Figure 7J, K). Furthermore, overexpression and knockdown of *NDRG1* in AML-12 cells revealed that NDRG1 significantly suppressed *GAMT* ubiquitination under CHX treatment (Figure 7L). Co-IP experiments further confirmed this result, and mutating *NDRG1* substantially increased GAMT ubiquitination (Figure 7M, N).

In summary, NDRG1 directly binds to GAMT through its critical residues Tyr-112, Ala-230, Ser-233, and Glu-238, thereby regulating GAMT ubiquitination and downstream creatine metabolism.

### 3.8 Creatine suppresses APAP-induced M1 polarization of macrophages through desuccinylation of PHB1 at Lys128

Previous studies have reported that succinylation plays an important role in the pathogenesis of APAP-induced liver injury(Yu, et al., 2024). By western blotting, we found that the overall succinylation level in macrophages from *NDRG1*^Hep-KO^ mice was significantly increased compared with that from *NDRG1*^Hep-flox^ mice, whereas exogenous creatine supplementation markedly reversed this increase, suggesting an inhibitory effect of creatine on macrophage succinylation (Figure 8A). This desuccinylation effect of creatine was further confirmed by global succinylation assays in macrophages isolated from creatine-treated mice (Figure 8B). Transcriptome sequencing revealed dynamic changes in succinylation-related enzymes, including *SIRT7*, *SIRT5*, *HAT1*, and *CPT1a*, in mouse macrophages before and after creatine treatment (Figure 8C). Accordingly, we performed succinylation site profiling on macrophages from creatine-treated and untreated mice and identified two sites, PHB1-K128 and PHB1-K208, that were significantly reduced in succinylation following creatine treatment (Figure 8F). Literature review indicated that PHB1 is a suppressor of macrophage M1 polarization and is closely associated with this process(Zhang, et al., 2025).

**Figure 8.**
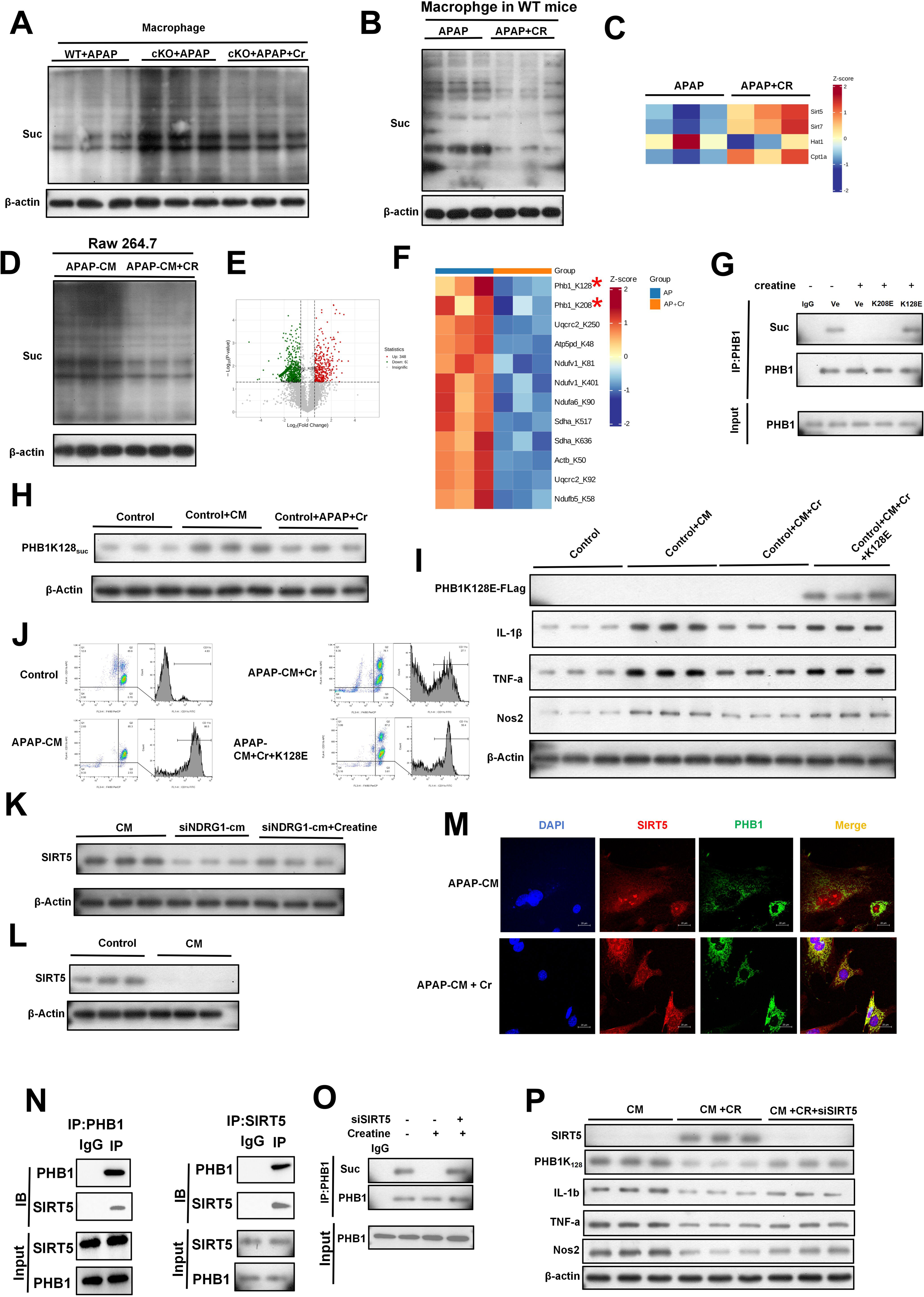
Creatine suppresses APAP-induced M1 polarization of macrophages through desuccinylation of PHB1 at Lys128 (A)Western blot detection of global succinylation levels in mouse liver macrophages.(B)Mouse liver macrophage Westernblot detection of global succinylation levels.*NDRG1*^Hep-flox^+APAP vs *NDRG1*^Hep-flox^ + APAP + creatine. (C) Mouse liver macrophage transcriptome sequencing heatmap.*NDRG1*^Hep-flox^+APAP vs *NDRG1*^Hep-flox^ + APAP + creatine.(D) Mouse macrophage Raw264.7 westernblot detection of global succinylation levels, CM=conditioned medium. (E) Mouse liver primary macrophage succinylation sequencing volcano plot.(F) Mouse liver primary macrophage succinylation sequencing heatmap.(G)Raw264.7 cell coip, IP:PHB1.(H)Western blot detection of Raw 264.7 PHB1-K128 site.(I)Western blot detection of Raw 264.7 Flag, IL-1β, TNF-α, NOS2, β-actin.(J)Raw 264.7 flow cytometry detection.(K)(L)Western blot detection of Raw 264.7 SIRT5 protein expression levels.(M)Raw264.7 immunofluorescence double staining, red=sirt5, green=PHB1.(N)Raw264.7 coip assay, left panel IP:PHB1, right panel IP:SIRT5.(O)Raw264.7 COIP assay. IP:PHB1. (P) Western blot detection of Raw 264.7 SIRT5, PHB1-K128, IL-1β, TNF-α, NOS2 protein expression levels.

Based on these findings, we generated constitutively succinylated mutant plasmids, *PHB1-K128E* and *PHB1-K208E*, and transfected them into RAW 264.7 cells. Co-immunoprecipitation (Co-IP) assays showed that creatine significantly reduced PHB1 succinylation (Figure 8G). Notably, the PHB1-K128E mutant, but not PHB1-K208E, reversed the creatine-induced decrease in succinylation, suggesting that PHB1-K128 may be one of the targets of creatine therapy (Figure 8G). Consistent with the Co-IP results, western blotting confirmed that succinylation at PHB1-K128 was markedly attenuated after creatine intervention (Figure 8H). Further experiments demonstrated that constitutive succinylation at PHB1-K128 significantly counteracted the therapeutic effect of creatine on M1 polarization, indicating that PHB1-K128 is a critical target through which creatine suppresses M1 polarization (Figure 8I, J). It has been previously reported that SIRT5 exerts a desuccinylation role in APAP-induced liver injury(Yu, et al., 2024). In our experiments, conditioned medium from NDRG1-knockdown AML12 cells significantly suppressed the expression of SIRT5 in macrophages, while creatine supplementation reversed this suppression (Figure 8K, L). Moreover, immunofluorescence co-localization suggested a potential interaction between PHB1 and SIRT5, which was further confirmed by Co-IP demonstrating a direct interaction (Figure 8M, N). Knockdown of SIRT5 significantly reversed the creatine-induced reduction in PHB1 succinylation and abolished the inhibitory effect of creatine on M1 polarization (Figure 8O, P).

Collectively, these findings indicate that creatine may exert its suppressive effect on macrophage M1 polarization by activating SIRT5-mediated desuccinylation of PHB1 at the K128 residue

### 3.9 Desuccinylation of PHB1 at K128 reverses the enhanced macrophage M1 polarization induced by hepatocyte-specific NDRG1 knockout

To further verify the role of PHB1-K128 in *NDRG1* cKO-mediated M1 polarization, we generated a PHB1-K128R desuccinylation mutant. Co-IP showed that K128R markedly reduced PHB1 succinylation (Figure 9A). Western blotting demonstrated that K128R significantly decreased M1 markers (IL-1β, NOS2, TNF-α) induced by si*NDRG1* AML12 conditioned medium, and flow cytometry and immunofluorescence confirmed attenuated M1 polarization in transfected RAW264.7 cells, indicating that PHB1-K128R partially reversed M1 polarization in vitro (Figure 9B, C, D).

**Figure 9.**
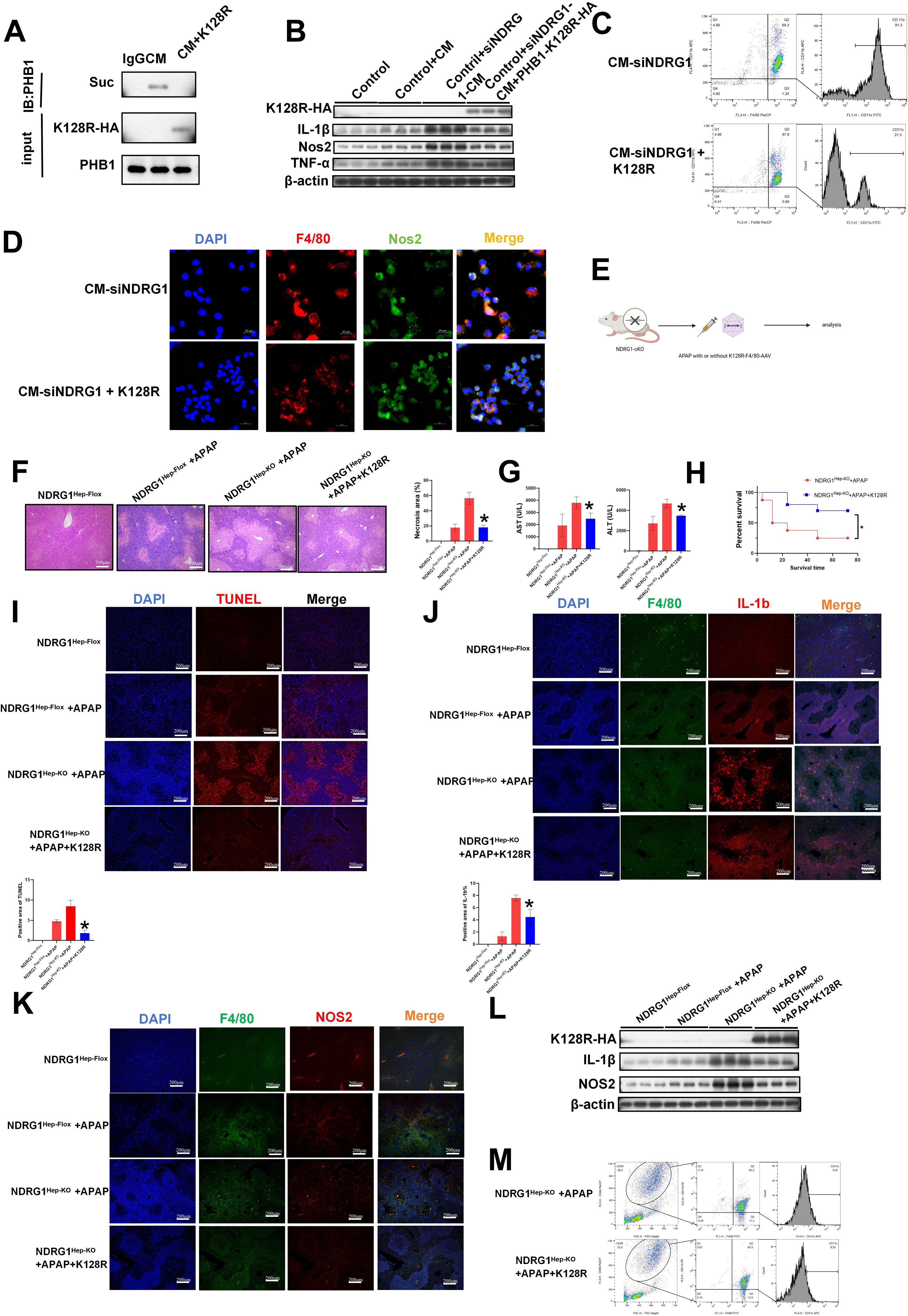
Desuccinylation of *PHB1* at K128 reverses the enhanced macrophage M1 polarization induced by hepatocyte-specific *NDRG1* knockout. (A) RAW264.7 co-IP assay, IP: PHB1. (B) Western blot detection of IL1-B, NOS2, and TNF-a expression in RAW 264.7 cells. (C) Flow cytometry detection of Raw 264.7 cells. (D) Immunofluorescence double staining detection of F4/80 and NOS2 expression. (E) Experimental design. (F) Mouse liver HE staining. Necrotic area, *, vs *NDRG1*^Hep-KO^+APAP, *P* < 0.05. (G) Mouse serum AST, ALT levels, *, vs *NDRG1*^Hep-KO^+APAP, *P* < 0.05. (H) Mouse K-M survival analysis, *P* < 0.05. (I) Mouse liver TUNEL staining, *, vs *NDRG1*^Hep-KO^+APAP, *P* < 0.05. (J) Mouse liver immunofluorescence double staining, green=F4/80, red=IL-1β, *, vs *NDRG1*^Hep-KO^+APAP, *P* < 0.05. (K) Mouse liver immunofluorescence double staining, green=F4/80, red=NOS2, *, vs *NDRG1*^Hep-KO^+APAP, *P* < 0.05. (L) Western blot detection of IL-1β and NOS2 protein expression levels. (M) Flow cytometry analysis of mouse liver non-parenchymal cells.

For in vivo validation, we constructed an AAV-F4/80 vector expressing PHB1-K128R for macrophage-specific mutation (Figure 9E). PHB1-K128R significantly ameliorated APAP-induced liver necrosis, transaminase elevation, hepatocyte apoptosis, and M1 polarization, and improved survival in *NDRG1*^Hep-KO^ mice (Figure 9F, G, H, I, J, K, L, M, *, vs *NDRG1*^Hep-KO^+APAP, *P* < 0.05). Collectively, PHB1-K128 is a key target mediating *NDRG1*-driven M1 polarization enhancement.

### 3.10 *PHB1*-K128 succinylation activates M1 polarization through the NF-κB pathway

To further investigate how PHB1 succinylation activates M1 polarization, we reviewed the literature and found that PHB1 is a negative regulator of the NF-κB pathway(Wang, et al., 2019; Jung, et al., 2020). Based on this finding, we examined NF-κB signaling by western blotting and found that the PHB1-K128E mutant promoted IκBα degradation and increased p65 expression (Supplementary Figure 2A). Moreover, immunofluorescence and western blotting revealed that PHB1-K128E significantly enhanced p65 nuclear translocation (Supplementary Figure 2B, C). Further experiments indicated that, upon PHB1-K128E mutation, the transcription of IL-1β and NOS2 was significantly upregulated compared with PHB1-oe, whereas treatment with an NF-κB inhibitor markedly suppressed these changes. This suggested that NF-κB might be a downstream transcription factor of PHB1-K128 succinylation (Supplementary Figure 2D, vs PHB1-oe, #, vs PHB1-K128E, *P* < 0.05). Subsequently, using a dual-luciferase reporter assay, we found that PHB1-K128E significantly activated NF-κB-driven transcription compared with PHB1-oe (Supplementary Figure 2E, *P* < 0.05). ChIP-PCR revealed that NF-κB could directly bind to the promoter sequences of iNOS and IL-1β, and the PHB1-K128E mutation significantly enhanced this binding relative to PHB1-oe (Supplementary Figure 2F, *P* < 0.05).Treatment with an NF-κB inhibitor significantly reversed the PHB1-K128E-induced aggravation of M1 polarization ((Supplementary Figure 2F). Collectively, these findings suggest that PHB1-K128 succinylation may activate NF-κB-mediated transcription of iNOS, IL-1β, and other genes by inhibiting the function of PHB1

### 3.11 *CXCL2*-positive macrophages may be the core cell population mediating M1 polarization in the context of hepatocyte-specific NDRG1 conditional knockout

Focusing solely on macrophage M1 polarization does not fully elucidate the role of *NDRG1* in APAP-induced liver injury. Based on our previous single-cell sequencing results, we found that the population of *CXCL2*-positive cells in *NDRG1* conditional knockout mice was significantly increased compared with *NDRG1*^Hep-flox^ mice following APAP-induced injury (Supplementary Figure 3A, B, C, D). Consistent with the single-cell sequencing data, western blotting, immunofluorescence, and flow cytometry all demonstrated that *CXCL2* expression in macrophages was markedly elevated after NDRG1 cKO (Supplementary Figure 3E, F, G). To validate the importance of this cell population, we constructed an AAV vector targeting macrophage-specific knockdown of *CXCL2*. Knockdown of *CXCL2* in macrophages significantly attenuated the increases in hepatic necrosis area, transaminase levels, and M1 polarization induced by *NDRG1* cKO (Supplementary Figure 3H, I, J, K, L. *, vs cKO+APAP, *P* < 0.05). These findings suggest that *CXCL2*-positive macrophages may be the target cell population mediating the protective role of *NDRG1*, and that intervention targeting *CXCL2* may be beneficial for alleviating APAP-induced pathology.

### 3.12 NDRG1 expression was verified using spatial transcriptomics and primary human hepatocytes-organoids (PHH-O)

Consistent with our results, spatial transcriptomic analysis of APAP-injured mouse and human livers showed that NDRG1 is primarily expressed in hepatocytes and markedly elevated upon APAP stimulation (Figure 10A, B). To verify the role of *NDRG1* in APAP-injured hepatocytes, human liver organoids were generated and challenged with APAP. At 100μM APAP, PHH-Os preserved basic morphology under light microscopy, and immunofluorescence revealed the most marked increase in NDRG1, consistent with earlier results (Figure 10C, E, F). The results also indicated that low concentrations of APAP significantly stimulated creatine production in human liver organoids, whereas high concentrations of APAP markedly inhibited creatine production, which might be attributable to extensive cell death in the organoids (Figure 10D). Consistent with the in vivo results, Western blotting revealed that NDRG1 protein expression was significantly increased in PHH-O organoids upon APAP stimulation (Figure 10G). In further organoid overexpression experiments, NDRG1 overexpression significantly upregulated GAMT expression (Figure 10I). Conditioned medium from NDRG1-overexpressing PHH-O organoids significantly inhibited macrophage polarization in hMDMs compared with control conditioned medium (Figure 10J). Moreover, exogenous creatine supplementation promoted PHB1 desuccinylation and suppressed macrophage polarization in hMDMs (Figure 10K, L). Together, these rescue and overexpression experiments demonstrated an NDRG1–creatine–succinylation circuit. Overall, spatial transcriptomics and PHH-O analyses both confirmed that NDRG1 was significantly upregulated in hepatocytes after APAP injury.

**Figure 10.**
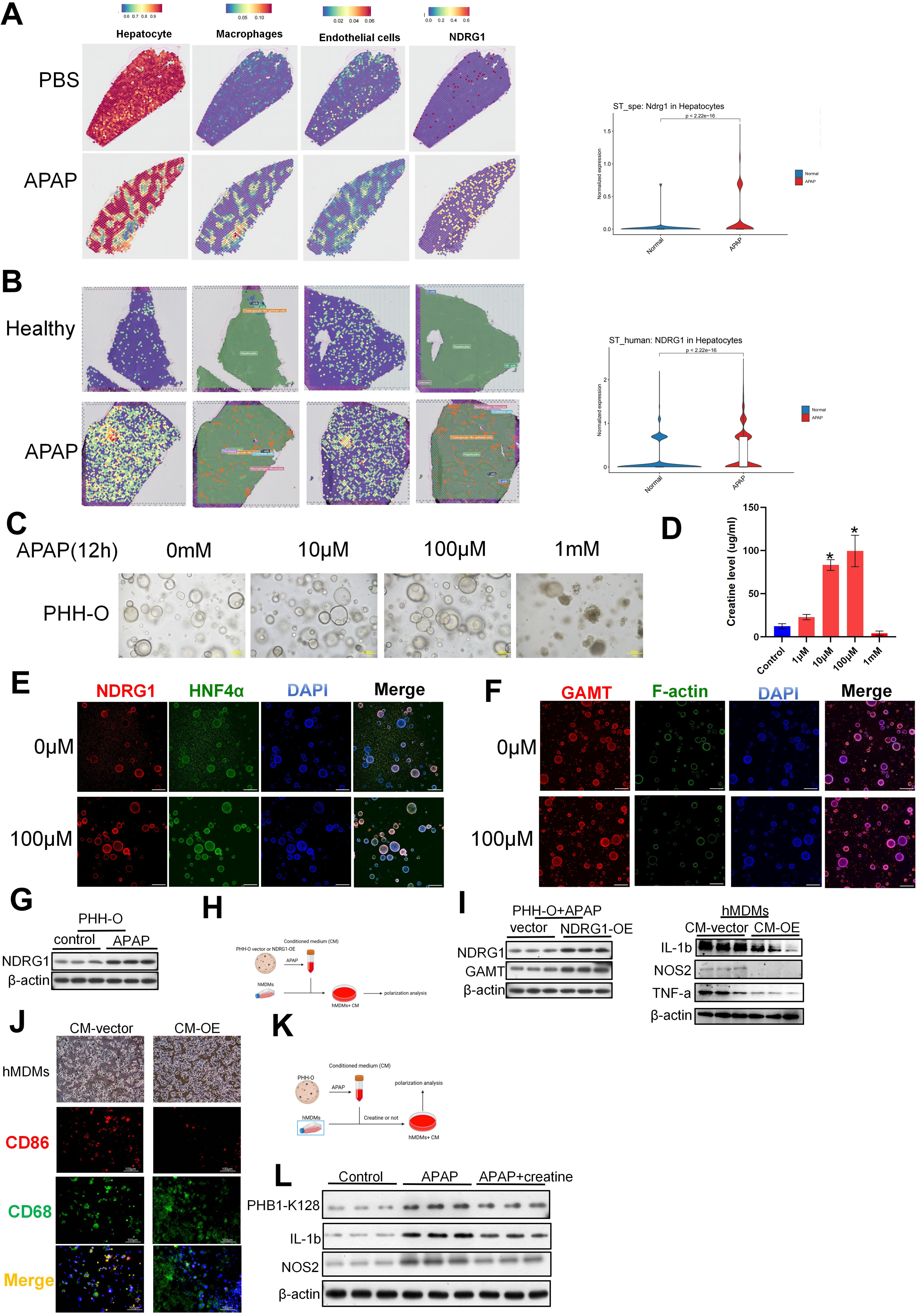
*NDRG1* expression was verified using spatial transcriptomics and human organ samples. (A) Spatial transcriptomic gene expression maps of mouse liver from dataset (PBS versus APAP).(B) Spatial transcriptomic gene expression maps of human liver from dataset GSE223559 (healthy versus liver failure). The expression of *NDRG1* is shown in these panels.(C) Morphological changes of human liver organoids following injury with different concentrations of APAP. Scale bar: 200 μm. (D) Creatine concentrations in the supernatant of organoids after injury with different concentrations of APAP (n = 3). (E) Co-staining of NDRG1 and HNF4A in human liver organoids after APAP injury. (F) Co-staining of GAMT and F-actin in human liver organoids after APAP injury. (G) Western blot analysis showing NDRG1 protein expression in PHH-O. (H) Schematic illustration of the experimental workflow for PHH-O. (I) Western blot analysis showing the protein expression of NDRG1, GAMT, IL-1β, TNF-α, and NOS2 in PHH-O and hMDMs. (J) Representative light micrographs and immunofluorescence double staining of hMDMs; red, CD86; green, CD68 (K) Schematic illustration of the experimental workflow. (L) Western blot analysis showing the protein expression of IL-1β, NOS2, and PHB1-K128 in hMDMs.

## 4. Discussion

In this study, we uncovered the protective roles of *NDRG1* and *GAMT* against APAP-induced liver injury and demonstrated that: 1) Hepatocyte-specific knockout of *NDRG1* significantly aggravates APAP-induced liver injury. 2) *NDRG1* in hepatocytes inhibits APAP-induced M1 polarization of macrophages by regulating creatine synthase and thus creatine production. 3) *NDRG1*-mediated creatine secreted by hepatocytes suppresses M1 macrophage polarization by promoting PHB1 desuccinylation. 4) Cxcl2⁺ macrophages may be one of the key cell populations mediating the protective effect of *NDRG1*.

There are several significant observations of this study. First, this study potentially represents the first report identifying a role for *NDRG1* in DILI. Previous studies have established NDRG1 as a stress-responsive gene that enhances cellular resistance to radiation and chemotherapy(Jung, et al., 2010; Yu, et al., 2024; Li, et al., 2025). As research has progressed, its protective function in acute organ injury has been increasingly recognized. Yuan et al. demonstrated that *NDRG1* significantly reduces reactive oxygen species (ROS) production during cardiac hypertrophy, thereby inhibiting its progression(Yuan, et al., 2025). Similarly, Zhang et al. found that inhibition of *NDRG1* exacerbates paraquat-induced lung injury, accompanied by significantly elevated levels of inflammatory cytokines, including IL-6, TNF-α, and IL-1β, suggesting an anti-inflammatory role for *NDRG1*(Zhang, et al., 2025). Consistent with these findings, our study reveals a significant protective effect of *NDRG1* against acetaminophen (APAP)-induced liver injury. We observed a marked increase in *NDRG1* expression in hepatocyte following APAP injury, indicating that *NDRG1* may function either as a protective gene or potentially as a damage-associated factor. Subsequent validation experiments demonstrated that, on the one hand, *NDRG1*-cKO mice exhibited significantly increased necrotic area and elevated inflammatory cytokine expression post-APAP injury compared to *NDRG1*^Hep-flox^ mice. On the other hand, conditioned medium from hepatocytes overexpressing *NDRG1* markedly suppressed the APAP-induced pro-inflammatory polarization of macrophages. These findings lay the groundwork for future investigations into the role of *NDRG1* in liver injury.

Second, with the advancement in understanding human metabolites and metabolomics sequencing technology, utilizing endogenous active metabolites for treating injury-related diseases has emerged as a promising therapeutic strategy for acetaminophen (APAP)-induced liver injury. Creatine, an amino acid containing a high-energy phosphate bond, is primarily synthesized in the liver(Kreider and Stout, 2021). Its production is tightly regulated by two key enzymes: L-arginine:glycine amidinotransferase (AGAT) and guanidinoacetate N-methyltransferase (GAMT)(Wyss and Kaddurah-Daouk, 2000). Previous studies have demonstrated that creatine not only promotes muscle synthesis and enhances exercise performance but also exerts therapeutic effects against liver and neurological damage(Barcelos, et al., 2016; Forbes, et al., 2022; Amiri and Sheikholeslami-Vatani, 2023; Desai, et al., 2025). Exogenous creatine supplementation has been shown to alleviate non-alcoholic fatty liver disease progression and doxorubicin-associated liver injury(Deminice, et al., 2015; Aljobaily, et al., 2020). Our study revealed that hepatic creatine levels were compensatorily elevated in mice following APAP challenge, accompanied by increased expression of GAMT. However, *NDRG1* knockout abolished this compensatory response, manifesting as reduced GAMT expression and diminished creatine levels. Further experiments demonstrated that *NDRG1* directly interacts with *GAMT* through four specific amino acid residues, thereby inhibiting GAMT ubiquitination. These findings identify *GAMT* and its downstream product creatine as critical therapeutic targets in our study. Nevertheless, neither creatine nor GAMT alone fully accounts for the protective role of *NDRG1*, warranting further in-depth investigations.

Third, the importance of succinylation is strongly demonstrated in this study. In previous studies, PHB1 has been recognized as a negative regulator of NF-κB and macrophage M1 polarization, and overexpression of PHB1 was found to suppress M1 macrophage polarization(Wang, et al., 2019). Succinylation is considered a post-translational modification that leads to loss of protein function. Consistent with these findings, we observed that hepatocyte-specific *NDRG1*-knockout mice exhibited increased global succinylation levels after modeling compared with *NDRG1*^Hep-flox^ mice, and succinylation sequencing also indicated that PHB1 succinylation was markedly reduced following creatine treatment. This suggests that PHB1 succinylation plays an important role in the protective effect mediated by NDRG1-induced creatine secretion. Further mechanistic exploration revealed that PHB1 succinylation may affect P65 nuclear entry. In summary, creatine was identified in this study as an agonist of SIRT5, exerting its desuccinylation effect by activating the desuccinylase.

Our study has several limitations. First, the clinical application of *NDRG1* agonists is not imminent, and the clinical utility of creatine remains largely confined to neurological disorders and sports-related injuries. Second, our investigation focused exclusively on the interaction between *NDRG*1 in hepatocytes and macrophages; its effects on other immune cell types were not explored. These aspects warrant further investigation in future studies.

In conclusion, our findings demonstrate that NDRG1 plays a critical protective role in APAP-induced liver injury. Mechanistically, NDRG1 modulates creatine production by affecting the ubiquitination of GAMT, the rate-limiting enzyme of creatine synthesis. The creatine regulated by NDRG1 in turn suppresses M1 polarization of liver macrophages via inhibiting succinylation of PHB1 at lysine 128.

## Abbreviations

APAP: Acetaminophen
NDRG1: N-myc downstream regulated gene-1
WT: Wild-type
RNA-Seq: RNA sequencing
BMDMs: bone marrow-derived macrophages
DILI: Drug-Induced Liver Injury
FDA: Food and Drug Administration
COVID-19: Coronavirus Disease 2019
AGAT: L-arginine:glycine amidinotransferase
GAMT: guanidinoacetate N-methyltransferase
cKO: conditional knockout
i.p.: intraperitoneal
PBS: Phosphate-Buffered Saline
ATCC: American Type Culture Collection
DMEM: Dulbecco’s Modified Eagle Medium
FBS: Fetal Bovine Serum
CM: conditioned medium
UMAP: Uniform Manifold Approximation and Projection
qPCR: quantitative Polymerase Chain Reaction
MPO: Myeloperoxidase
TUNEL: Terminal deoxynucleotidyl transferase dUTP Nick-End Labeling
GSEA: Gene Set Enrichment Analysis
IL: Interleukin
TNF: Tumor Necrosis Factor
NOS2: Nitric Oxide Synthase 2
iNOS: inducible Nitric Oxide Synthase
FACS: Fluorescence-Activated Cell Sorting
KEGG: Kyoto Encyclopedia of Genes and Genomes
Co-IP: Co-Immunoprecipitation
PHB1: Prohibitin 1
SIRT5: Sirtuin 5
SIRT7: Sirtuin 7
HAT1: Histone Acetyltransferase 1
CPT1a: Carnitine Palmitoyltransferase 1A
NF-κB: Nuclear Factor kappa-light-chain-enhancer of activated B cells
IκBα: Inhibitor of NF-κB alpha
ChIP: Chromatin Immunoprecipitation
CXCL2: C-X-C Motif Chemokine Ligand 2
AAV: Adeno-Associated Virus
PHH-O: Primary Human Hepatocytes-Organoids
ROS: Reactive Oxygen Species
GEO: Gene Expression Omnibus
NSFC: National Natural Science Foundation of China
ARRB1: Arrestin Beta 1
eIF2α: Eukaryotic Initiation Factor 2 Alpha
ER: Endoplasmic Reticulum
LPLUNC1: Long Palate, Lung and Nasal Epithelium Carcinoma Associated Protein 1
TRIM21: Tripartite Motif Containing 21
KAT2A: Lysine Acetyltransferase 2A
ALDH2: Aldehyde Dehydrogenase 2
MASLD: Metabolic dysfunction-associated Steatotic Liver Disease
cGAS-STING: cyclic GMP-AMP Synthase-Stimulator of Interferon Genes

## Author contribution

Y. J was the as the submission’s guarantor. Y. J. and J.Y designed and conducted trials and analyzed the data. C.H. and C.Y. and Y.H. made figures and tables and wrote the manuscript. Y.C performed the blinded allocation of the animals. H.C and H. X. supervised the project, designed the experiments, helped with the data interpretation, participated in the data analysis, and wrote the manuscript. Dr. Felix W. Leung contributed to the conception and design of the study, supervised all aspects of the research, and was critically involved in the interpretation of data and the revision of the manuscript for important intellectual content. All authors approved the final version of the article.

## Data availability

The datasets used and/or analyzed during the current study are available from the corresponding author on reasonable request.

## Acknowledgements

We thank Professor Felix W Leung at University of California Los Angeles David Geffen School of Medicine for providing the guidance. We thank the Gene Expression Omnibus (GEO) for providing the sequencing data.

---

**Supplementary Figure 1.**
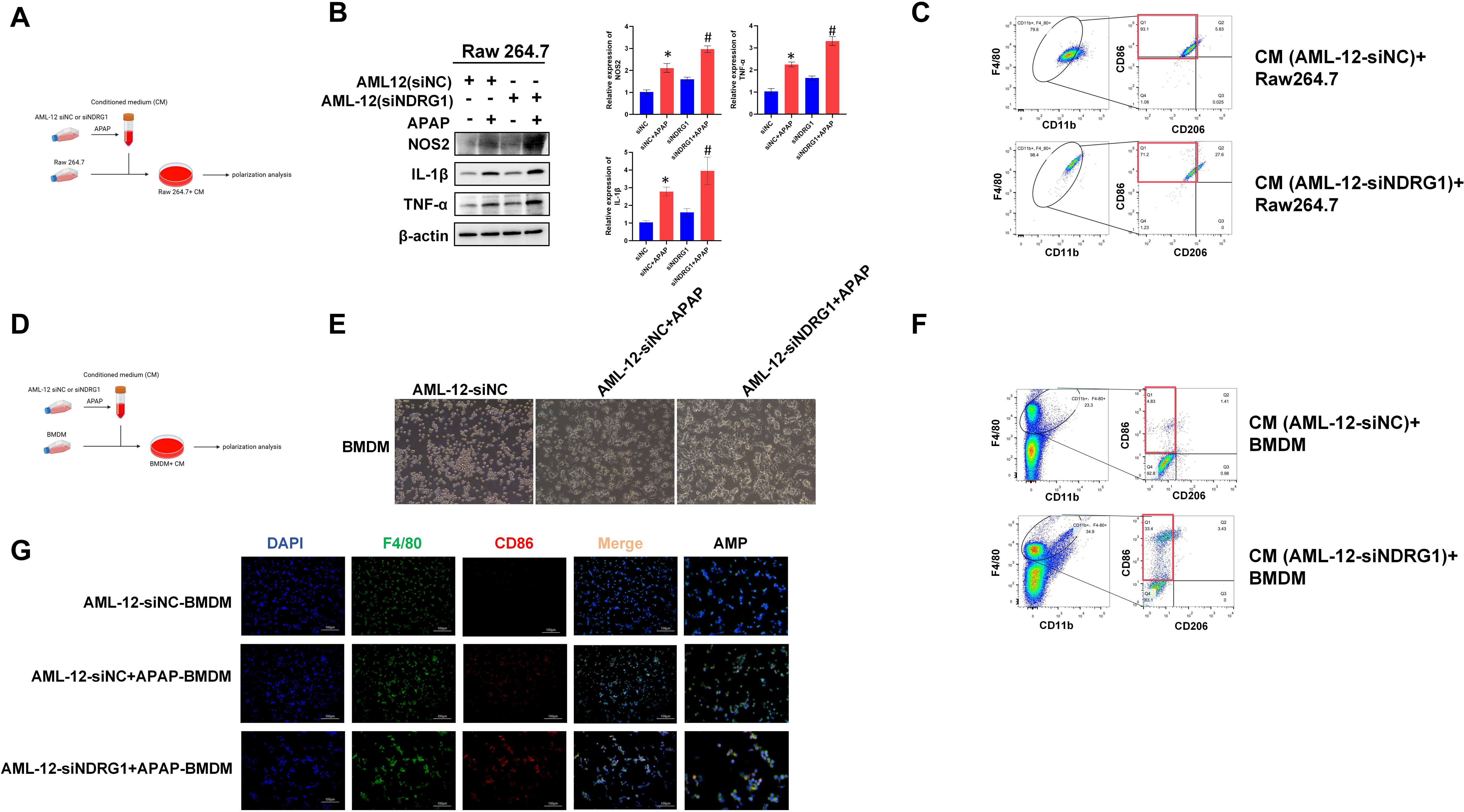
(A) Schematic diagram of *in vitro* study. RAW 264.7 were cultured with conditioned medium from AML-12 pre-activated by APAP. (B) Western blot of M1 macrophage polarization markers in RAW 264.7. * vs siNC *P* < 0.05, # vs siNC + APAP *P* < 0.05. n=3. (C) Flow cytometric analysis of RAW 264.7 (F4/80^+^, CD86^+^, CD11b^+^). (D) Schematic diagram of *in vitro* study. BMDM were cultured with conditioned medium from AML-12 pre-activated by APAP. (E) Light microscope of BMDM. (F) Flow cytometric analysis of BMDM (F4/80^+^, CD86^+^, CD11b^+^). (G) Dual immunofluorescence staining of F4/80 and CD86 in BMDM.

**Supplementary Figure 2.**
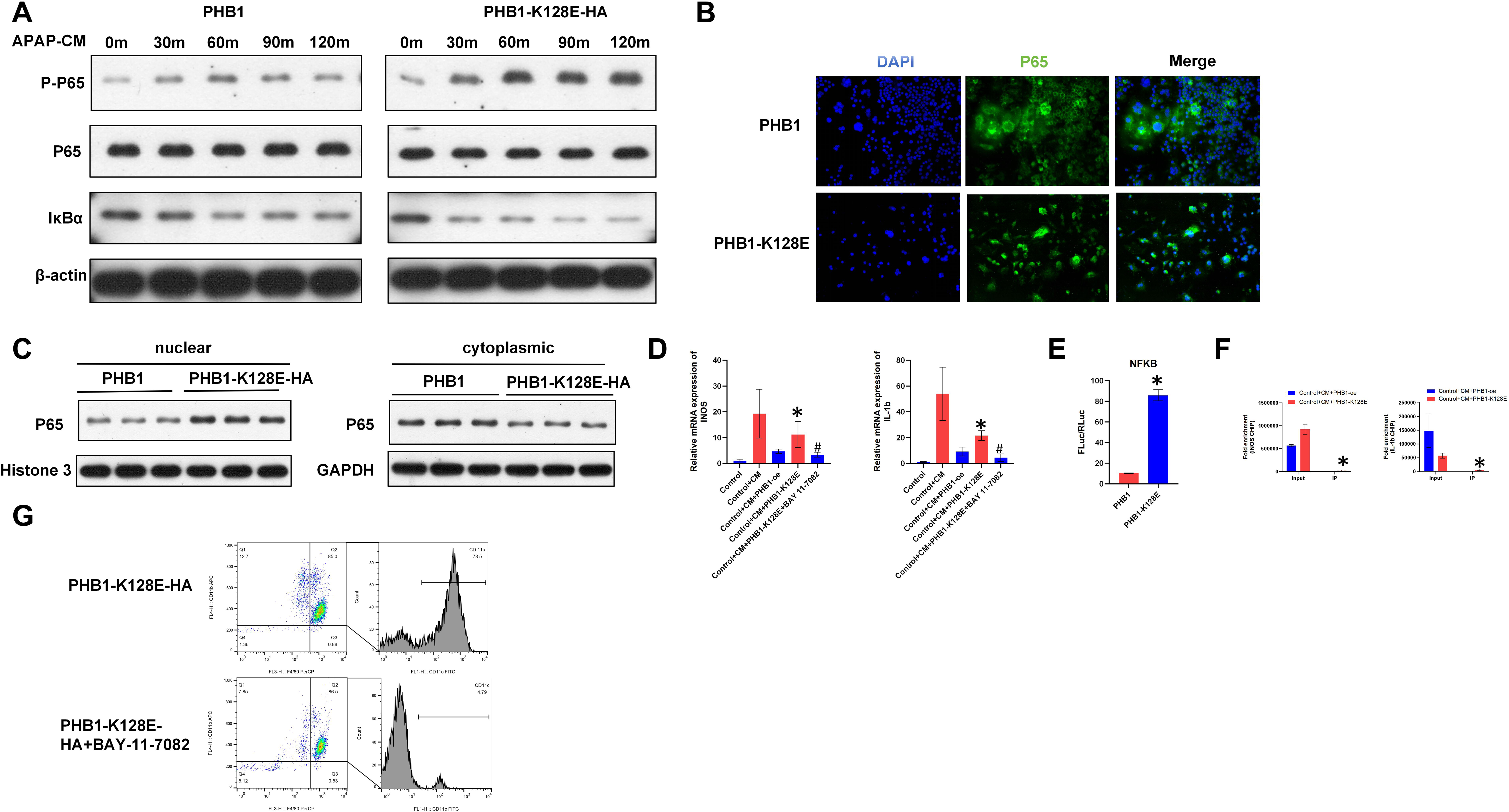
(A) Western blot detection of PP65, p65, and IκBα expression in RAW 264.7 cells. Vector vs PHB1-K128E-HA. (B) P65 immunofluorescence staining. (C) Western blot detection of nuclear and cytoplasmic P65 protein expression. (D) qPCR analysis of IL-1β and NOS2 mRNA levels in RAW264.7 cells. vs PHB1-oe, #, vs PHB1-K128E, *P* < 0.05. (E) NF-κB luciferase reporter assay, *, *P* < 0.05. (F) ChIP-qPCR detection of IL-1β and NOS2 transcription, *, *P* < 0.05. (G) Flow cytometry analysis of RAW 264.7 cells.

**Supplementary Figure 3.**
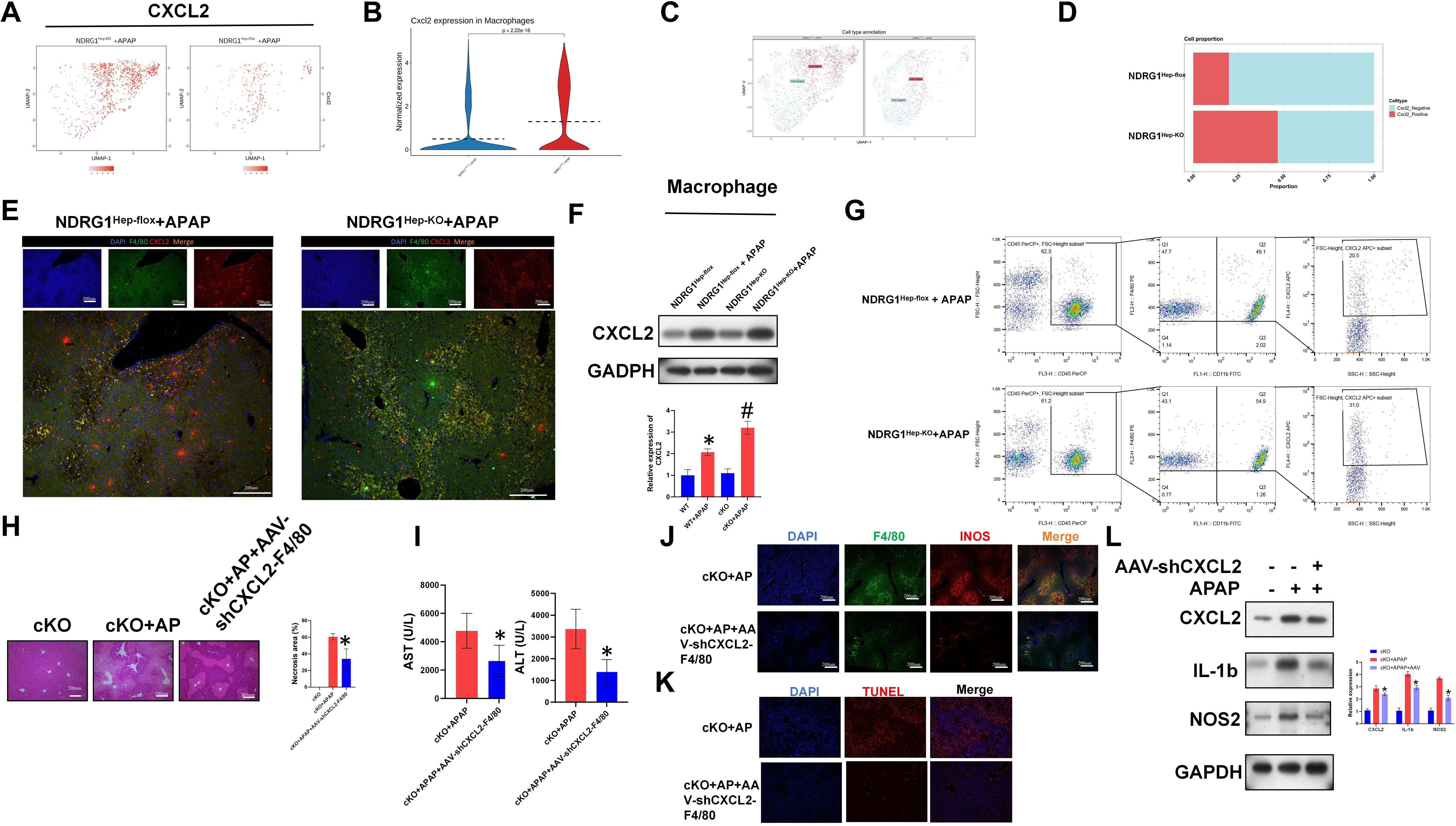
*CXCL2*-positive macrophages may be the core cell population mediating M1 polarization in the context of hepatocyte-specific *NDRG1* conditional knockout (A) Feature plot of CXCL2 in macrophages, *NDRG1*^Hep-KO^ vs *NDRG1*^Hep-flox^. (B) Violin plot showing differential expression of CXCL2 in macrophages. (C) UMap plot showing distribution of CXCL2-positive and CXCL2-negative macrophages in *NDRG1*^Hep-flox^ and *NDRG1*^Hep-KO^. (D) Proportion of CXCL2-positive macrophages in *NDRG1*^Hep-flox^ and *NDRG1*^Hep-KO^ mice. (E) Immunofluorescence double staining in mouse liver, green=F4/80, red=CXCL2. (F) Western blot revealing CXCL2 expression in mouse liver macrophages, *, vs *NDRG1*^Hep-flox^ #, vs *NDRG1*^Hep-flox^+APAP, *P* < 0.05. (G) Flow cytometry analysis of mouse liver non-parenchymal cells. (H) Mouse liver HE staining, necrotic area, *, vs *NDRG1*^Hep-KO^+APAP, *P* < 0.05. (I) Mouse serum AST, ALT levels, *, vs *NDRG1*^Hep-KO^+APAP, *P* < 0.05. (J) Mouse liver immunofluorescence double staining, green=F4/80, red=INOS, *, vs *NDRG1*^Hep-KO^+APAP, *P* < 0.05. (K) Mouse liver TUNEL staining, *, vs *NDRG1*^Hep-KO^+APAP, *P* < 0.05. (L) Western blot detection of CXCL2, IL-1β, and NOS2 expression in mouse liver macrophages, *, vs *NDRG1*^Hep-KO^+APAP, *P* < 0.05.

